# RNA structure probing with 4-base-recognizing RNA cutters

**DOI:** 10.64898/2026.11.27.769052

**Authors:** Masayuki Takahashi, Masayuki Nashimoto

## Abstract

To elucidate dynamic higher-order structures of RNA in human cells would be important not only to deeply understand the physiological functions of mRNA and non-coding RNA but also to establish precision medicine. tRNase Z^L^ together with hook-type sgRNA works as a 4-base-recognizing RNA cutter. We thought that RNA structure probing using the RNA cutters with hook-type sgRNAs of various 4-base recognition sequences would provide valuable information on the higher-order structures of cellular RNA. In this study, we investigated the feasibility of this method *in vitro*. We examined nine 32-nt RNA targets of various degrees of folding for cleavability by human tRNase Z^L^ guided by each of thirteen hook-type sgRNAs. The assays were carried out under four different conditions: at 37 or 50°C with MgCl_2_ or spermidine. From these assays, we identified four main findings: At least two consecutive G or C nt in the target RNA recognition sequence are needed for hook-type sgRNA to work as a guide for tRNase Z^L^. A longer target RNA recognition sequence of hook-type sgRNA provides better guiding ability, and the hook-type sgRNA that can form with target RNA a hairpin structure without bulged nucleotides shows a better guiding ability. tRNase Z^L^ with hook-type sgRNA, which was designed to work as a 4-base-recognizing RNA cutter, can also work efficiently as a 3-base-recognizing RNA cutter in some cases. Cleavage of target RNA by tRNase Z^L^ guided by hook-type sgRNA occurs only in unstructured or loosely-folded target RNA, suggesting that sgRNA-guided tRNase Z^L^ can discriminate less folded regions from tightly folded ones in long cellular mRNA or non-coding RNA. Structure probing of cellular RNA by introducing into the cells the hook-type sgRNA designed based on the findings of this study would provide valuable information on the higher-order structures of the cellular RNA.

## Introduction

To elucidate dynamic higher-order structures of RNA in human cells would be important not only to deeply understand the physiological functions of mRNA and non-coding RNA but also to establish precision medicine [1,2]. Although prediction of the cellular RNA structures for these purposes would become possible with artificial intelligence and/or quantum computers in the future, we still need to perform various types of experiments, which would be in turn used to improve the computer prediction [3–6]. The RNA structure probing experiments include X-ray crystallography, nuclear magnetic resonance spectroscopy, cryogenic electron microscopy, enzymatic probing with RNases such as RNase A, RNase T1 and RNase V1, and chemical probing with chemical reagents such as dimethyl sulfate, 2-methylnicotinic acid imidazolide and 1-ethyl-3-(3-dimethylaminopropyl)carbodiimide [7–9]. The last one is also applicable to *in vivo* experiments. In the development of oligonucleotide therapeutics such as conventional antisense oligonucleotides (ASOs) and small interfering RNAs (siRNAs), the structure probing of target RNA would be useful to select candidate target sites and to streamline the screening process.

This would also hold in the development of small guide RNA (sgRNA) for tRNase Z^L^-utilizing efficacious (TRUE) gene silencing [10]. TRUE gene silencing is a potential gene suppression method that we have been trying to put into practical use. The principle of this method is that target RNA can be cleaved by the tRNA 3′ processing endoribonuclease tRNase Z^L^ through forcing the target RNA to form a pre-tRNA-like or micro-pre-tRNA-like complex with a 7–30-nt sgRNA. The cleavage occurs immediately after a nucleotide corresponding to the discriminator of tRNA (D-nt) and/or 1/2-nt downstream of it. sgRNAs are categorized into 4 groups, heptamer type, linear type, hook type and 5′-half-tRNA type. Recently we have examined six 14-nt linear-type sgRNAs targeting six different sites of the human MALAT1 RNA, which were selected without using structure probing data, for their ability to reduce the RNA level in A549 cells [11]. And we have observed that only three of them show the reducing effect but only moderately. One of the strategies to improve the effect would definitely be to select target sites in much less folded RNA regions based on RNA structure probing data.

In the course of planning structure probing of the MALAT1 RNA, we came up with a new method for RNA structure probing based on the RNase 65 activity. Incidentally, TRUE gene silencing has been derived from the intensive study of the RNase 65 activity [10,12–15]. RNase 65 is a complex of tRNase Z^L^ and 3′-truncated tRNA that lacks 9 nt such as tRNA^Arg^ or tRNA^Ala^ and works as a 4-base-recognizing RNA cutter [15]. An RNA substrate is recognized by RNase 65 through four Watson-Crick-Franklin (WCF) base-pairings with the 3′-truncated tRNA. For example, the 3′-truncated tRNA^Arg^ recognizes the substrate via WCF base-pairings of its 5′-terminal sequence 5′-GGGC-3′ with the sequence 5′-GCCC-3′ of the substrate RNA. RNase 65 with the 3′-truncated tRNA^Arg^ and tRNA^Ala^ are named RNase 65R and RNase 65A, respectively.

We have observed that the 344-nt RNA mostly derived from mouse 18S rRNA can be cleaved only after the 111th nucleotide (D-nt) that is 1-nt downstream of the 5′-GCCC-3′ sequence by RNase 65R *in vitro*, although the RNA contains the 5′-GCCC-3′ sequence at three sites [16]. This observation suggests that RNase 65R can discriminate less folded regions from tightly folded ones in the target RNA. We have also demonstrated that tRNase Z^L^ works as a 4-base-recognizing RNA cutter together with ∼26-nt hook-type sgRNA instead of 3′-truncated tRNA, although tRNase Z^L^ may not form a complex with the hook-type sgRNA in the absence of its target RNA [17,18]. From these observations, we thought that the RNA structure probing using tRNase Z^L^ together with hook-type sgRNAs of various 4-base recognition sequences would provide further information on the higher-order structures of the target RNA. This method would also be applicable to cellular target RNA by introducing the hook-type sgRNA into the cells.

In this study, we investigated the feasibility of this method with short RNA targets of various degrees of folding. The nine 32-nt RNA targets were examined for their specific cleavability by tRNase Z^L^ together with each of the thirteen hook-type sgRNAs *in vitro*. We showed that cleavage of target RNA by tRNase Z^L^ guided by hook-type sgRNA occurs only in unstructured or loosely-folded target RNA, and we believe that this method could be used to obtain additional information on the higher-order structures of RNA.

## Materials and Methods

### Target RNA synthesis

Nine 32-nt 5′-6-carboxyfluorescein (FAM)-labeled RNA targets, Target-1 (5′-GAAAAAACCCCAAAAAAGCCCAAAAAAAAAAA-3′), Target-2 (5′-GAAAAGAGAACCAAAAAAGUACAUAUAAAAAA-3′), Target-3 (5′-GAAAAGAGAACCAAAAAGCGCGAUUUUAAAAA-3′), Target-4 (5′-GAAAAAAGGGGAAAAAACGGGAAAAAAAAAAA-3′), Target-5 (5′-GGACCCCAGCCCUAAAAAAAGGGCUGGGGUCC-3′), Target-6 (5′-GGAAUCUAGCCCGAGACCCCUAGCGCAGUUGU-3′), Target-7 (5′-GAGGGAGAUUUGUUUUAUAUGUGUCUAACCUU-3′), Target-8 (5′-UGGCCGUGCGGGGUUGUACAACCGAGAUUACU-3′) and Target-9 (5′-GGAAUCUGGCCCGAGGCCCCUGGCGCAGUUGU-3′) were chemically synthesized and subsequently purified by high-performance liquid chromatography by Nippon Bioservice (Saitama, Japan).

### RNA secondary structure prediction

Potential RNA secondary structures of Target-1–Target-9 were calculated on the website http://rna.tbi.univie.ac.at/cgi-bin/RNAWebSuite/RNAfold.cgi.

### Hook-type sgRNA synthesis

The following 13 hook-type sgRNAs, which were 5′-phosphorylated and fully 2′-*O*-methylated with full phosphorothioate linkages, were synthesized by Nippon Bioservice as above: Hook-1 (5′-pGGGCCAGCCAGGUUCGACUCCUGGCUG-3′), Hook-2 (5′-pGGGCCAGCCAGGUUCGACUCCUGGCU-3′), Hook-3 (5′-pGGGGCAGCCAGGUUCGACUCCUGGCU-3′), Hook-3.1 (5′-pGGGGCAGCCAGGUUCGACUCCUGGCUG-3′), Hook-4 (5′-pGCGCCAGCCAGGUUCGACUCCUGGCU-3′), Hook-5 (5′-pAAAUCAGCCAGGUUCGACUCCUGGCU-3′), Hook-6 (5′-pAAAACAGCCAGGUUCGACUCCUGGCU-3′), Hook-7 (5′-pAUAUCAGCCAGGUUCGACUCCUGGCUG-3′), Hook-8 (5′-pGUACCAGCCAGGUUCGACUCCUGGCUG-3′), Hook-9 (5′-pGGUUCAGCCAGGUUCGACUCCUGGCUG-3′), Hook-10 (5′-pUCUCCAGCCAGGUUCGACUCCUGGCUG-3′), Hook-11 (5′-pCCCCCAGCCAGGUUCGACUCCUGGCUG-3′) and Hook-12 (5′-pCCCGCAGCCAGGUUCGACUCCUGGCU-3′).

### Preparation of recombinant human tRNase Z^L^

The histidine-tagged human tRNase Z^L^ without its N-terminal 30 amino acids was expressed using the expression plasmid pQE-80L in *E. coli* strain Rosetta(DE3)pLysS and purified with nickel-agarose beads as before [19].

### *In vitro* RNA cleavage assay

*In vitro* RNA cleavage assay was performed at 37 or 50°C for 30 min with 1 pmol of 5′-FAM-labeled target RNA, 1 pmol of hook-type sgRNA and 8 pmol of recombinant human tRNase Z^L^ in a 20-μl mixture containing 10 mM Tris-HCl (pH 7.1) and 5 mM MgCl_2_ or 0.2 mM spermidine. Reaction products were resolved on a 20% polyacrylamide gel containing 8 M urea and analyzed using a Gel Doc EX imager (Bio-Rad, Tokyo, Japan).

## Results

### Target RNAs and hook-type sgRNAs for tRNase Z^L^

We designed nine 32-nt RNA targets, Target-1–Target-9, and chemically synthesized them with a FAM moiety at their 5′ terminus (Fig 1). Target-1–Target-4 were designed to avoid forming secondary structures, whereas Target-5 was designed to tightly fold with the Gibbs free energy change (△G) = -29.2 kcal/mol. Target-6–Target-9 were designed to loosely fold with △G = -2.5–5.8 kcal/mol. Each RNA target was designed to have multiple target sequences of the hook-type sgRNAs.

**Fig 1.**
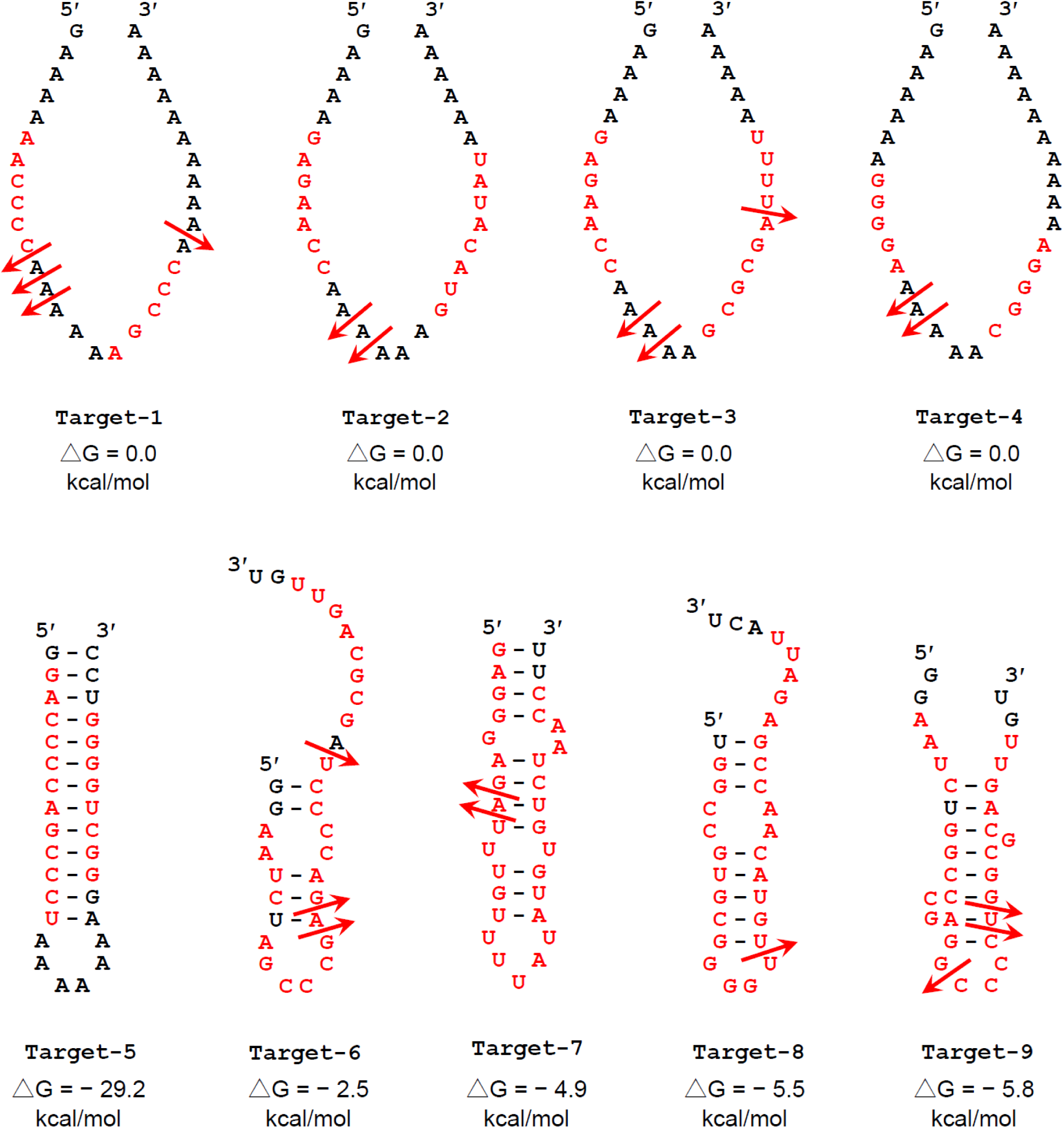
Predicted secondary structures of RNA targets, Target-1–Target-9, with △G values. Bases potentially involved in binding of at least one of the hook-type sgRNAs Hook-1–Hook-12 are shown in red. An arrow indicates the position where cleavage by tRNase Z^L^ occurred under at least one of the four assay conditions.

Thirteen hook-type sgRNAs were designed and chemically synthesized with modifications of a 5′-phosphate, full 2′-*O*-methylation and full phosphorothioate linkages (Fig 2). The reason why we synthesized the hook-type sgRNAs with these chemical modifications is to confirm that these modifications do not inhibit the guiding ability of the hook-type sgRNAs, in this *in vitro* study, because these modifications, which are thought to extend RNA’s lifetime in human cells, should be needed for future *in vivo* experiments. Hook-1 and Hook-2 were designed to target the 5′-GCCC-3′ sequence, and Hook-3 and Hook-3.1 were to target the 5′-CCCC-3′ sequence. The sgRNAs Hook-4 to Hook-12 were designed to target the 5′-GCGC-3′, 5′-AUUU-3′, 5′-UUUU-3′, 5′-AUAU-3′, 5′-GUAC-3′, 5′-AACC-3′, 5′-GAGA-3′, 5′-GGGG-3′ and 5′-CGGG-3′ sequences, respectively. Considering the non-WCF G-U base pair, these sgRNAs can target more sequences. Hook-2, Hook-3, Hook-4, Hook-5, Hook-6, and Hook-12 can also recognize the target RNA by base-pairings with the 2nd to 5th nt sequence or the 1st to 5th nt sequence of the hook-type sgRNAs. Hook-2 and Hook-3 have a one-base-pair shorter stem structure than Hook-1 and Hook-3.1, respectively. These were designed to examine how the one-base-pair difference affects the interaction between sgRNA and target RNA and the efficiency of target RNA cleavage by tRNase Z^L^.

**Fig 2.**
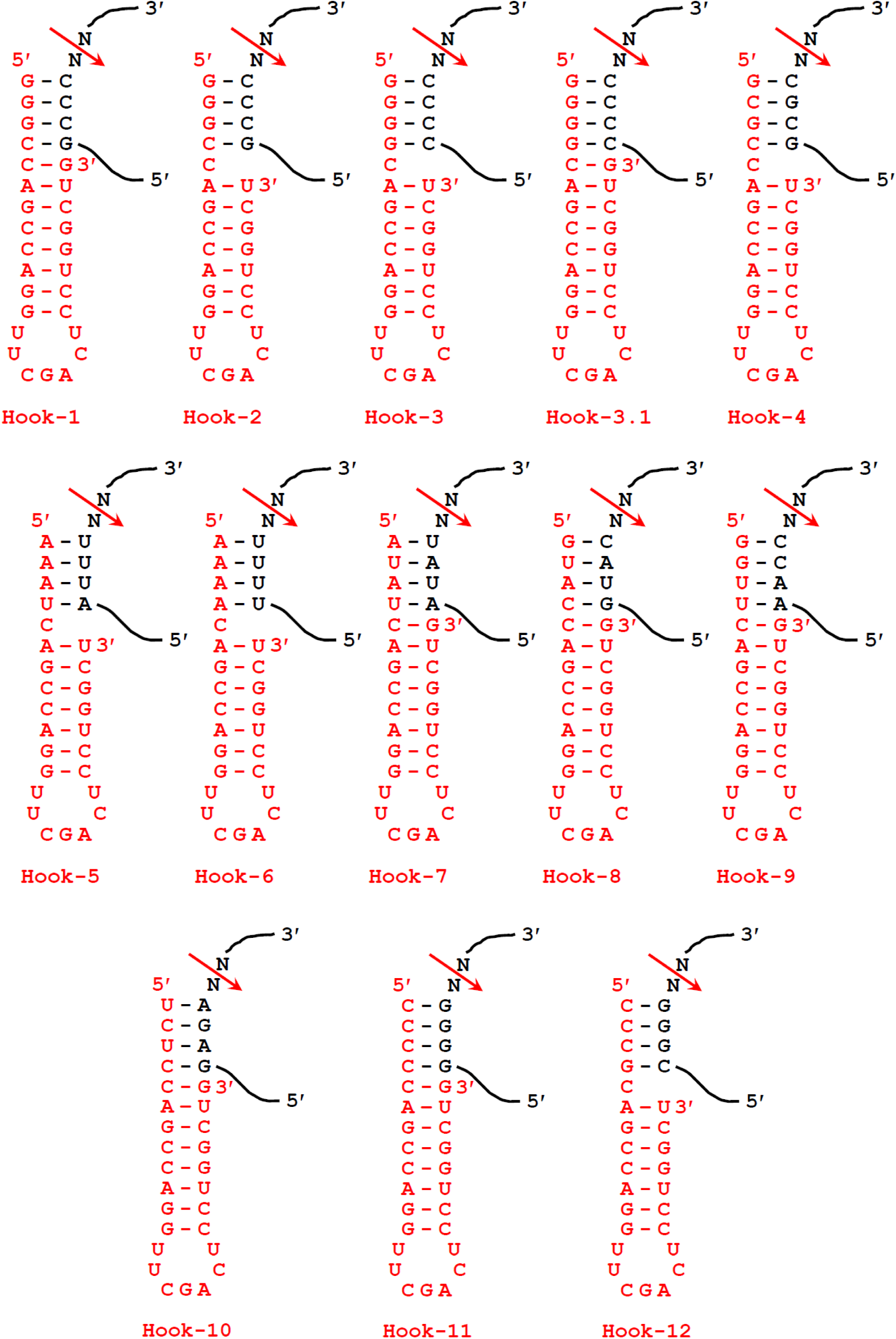
Potential secondary structures of complexes between target RNA and hook-type sgRNA. An arrow denotes the expected tRNase Z^L^ cleavage site. N’s are A, U, G or C.

### Cleavage assays for unstructured Target-1

Target-1 had four potential sgRNA binding sites: one for Hook-1, Hook-2 and Hook-3.1 and two for Hook-3 and Hook-9 (S1–S4 Fig. and S10 Fig.). Cleavage assays for Target-1 were performed at 37 and 50°C in the presence of 5 mM MgCl_2_ (Fig 3A) and 0.2 mM spermidine (Fig 3B), and cleavage sites were determined on a sequencing gel with an alkaline ladder as size standards (Fig 3C).

**Fig 3.**
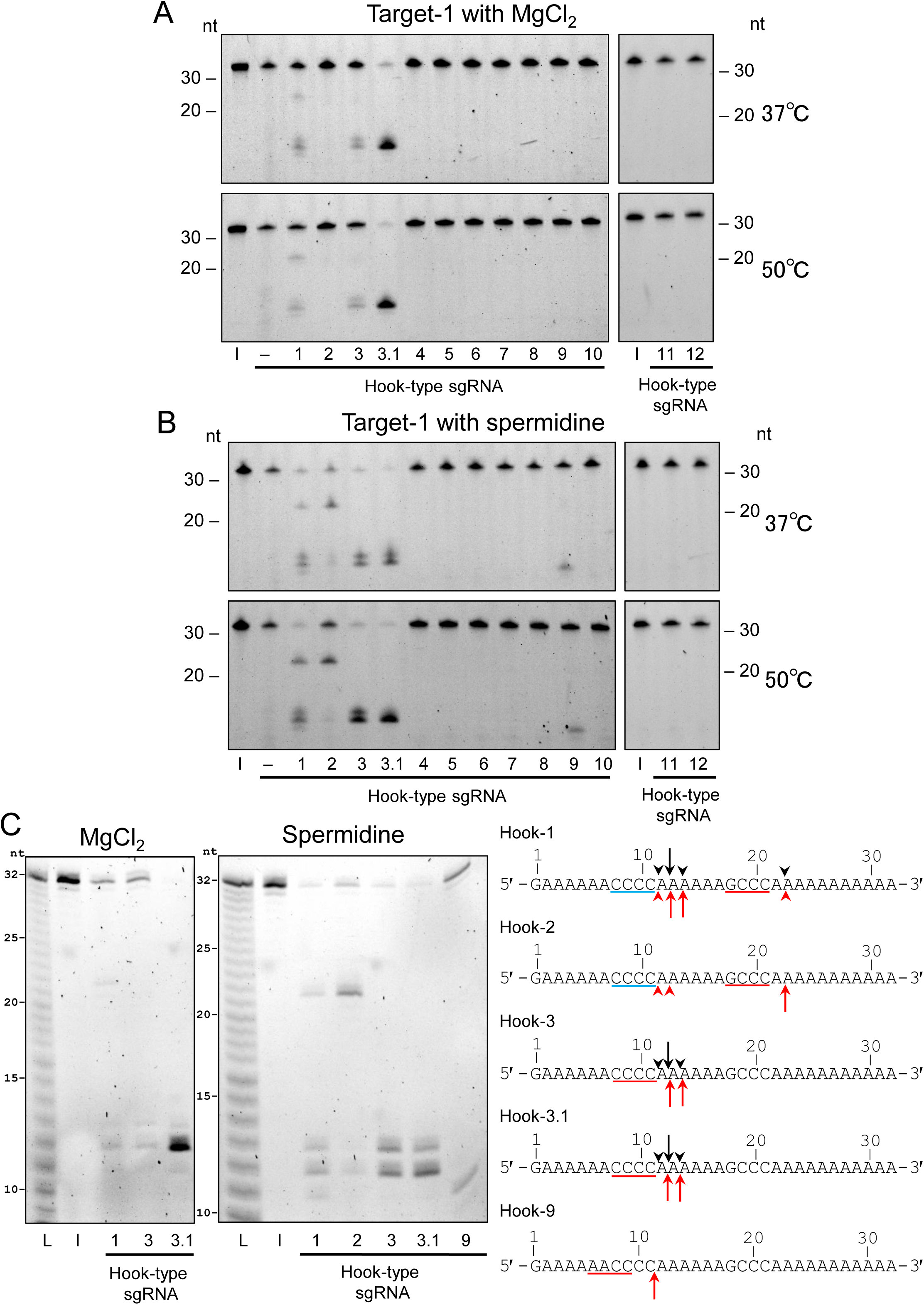
The tRNase Z^L^ cleavage assay for Target-1 in the presence of the hook-type sgRNAs Hook-1–Hook-12. The cleavage assays were performed with 5 mM MgCl_2_. (A) or 0.2 mM spermidine (B) at 37 or 50°C, and the reaction products were separated on a denaturing 12.5% polyacrylamide gel. (C) The cleaved RNA products in the reactions with MgCl_2_ at 50°C and those with spermidine at 37°C were also separated on a denaturing 20% polyacrylamide sequencing gel. The sites of cleavages with MgCl_2_ and with spermidine are indicated by black and red arrows/arrowheads, respectively, on the target RNA. A relatively strong cleavage site is denoted by an arrow. The potential sgRNA binding sequences of ≧ 4 nt and < 4 nt are indicated by red and blue lines, respectively. 20-and 30-nt RNAs and an alkaline ladder were used as size standards. L, alkaline ladder; I, input RNA.

Under any of those conditions, Target-1 was cleaved by Hook-1-guided tRNase Z^L^ after the 22nd nt A (D-nt) as expected, and, in addition, it was cleaved unexpectedly around the 12th nt A (Fig 3 and S1 Fig.). Hook-2-guided tRNase Z^L^ cleaved Target-1 in a similar fashion but only with spermidine (Fig 3 and S2 Fig.). The event of the cleavages around the 12th nt A suggests that three consecutive G-C WCF base-pairings are sufficient for Hook-1-and Hook-2-guided tRNase Z^L^ to recognize the target RNA.

Target-1 was cleaved around the 12th nt A (D-nt) by Hook-3-and Hook-3.1-guided tRNase Z^L^ in a similar way, although the latter cleaved the target more efficiently than the former in the presence of MgCl_2_ (Fig 3, S3 Fig. and S4 Fig.). Hook-9-guided tRNase Z^L^ cleaved Target-1 1-nt downstream of a D-nt (the 10th nt C) only with spermidine (Fig 3 and S10 Fig.).

### Cleavage assays for unstructured Target-2

Target-2 had also four potential sgRNA binding sites: one for Hook-7, Hook-8, Hook-9 and Hook-10 (S8–S11 Fig.). The following cleavage assays and cleavage site determination for the other RNA targets were performed in the same way as those for Target-1.

Target-2 was cleaved efficiently by Hook-9-guided tRNase Z^L^ 1/2-nt downstream of a D-nt (the 13th nt A) only with spermidine (Fig 4 and S10 Fig.). And, unusually, it was also cleaved very faintly at the same positions by Hook-1- and Hook-3.1-guided tRNase Z^L^ (Fig 4, S1 Fig. and S4 Fig.). This observation suggests that Hook-1- and Hook-3.1-guided tRNase Z^L^ can recognize target RNA by only two consecutive G-C WCF base-pairings (Fig 4C).

**Fig 4.**
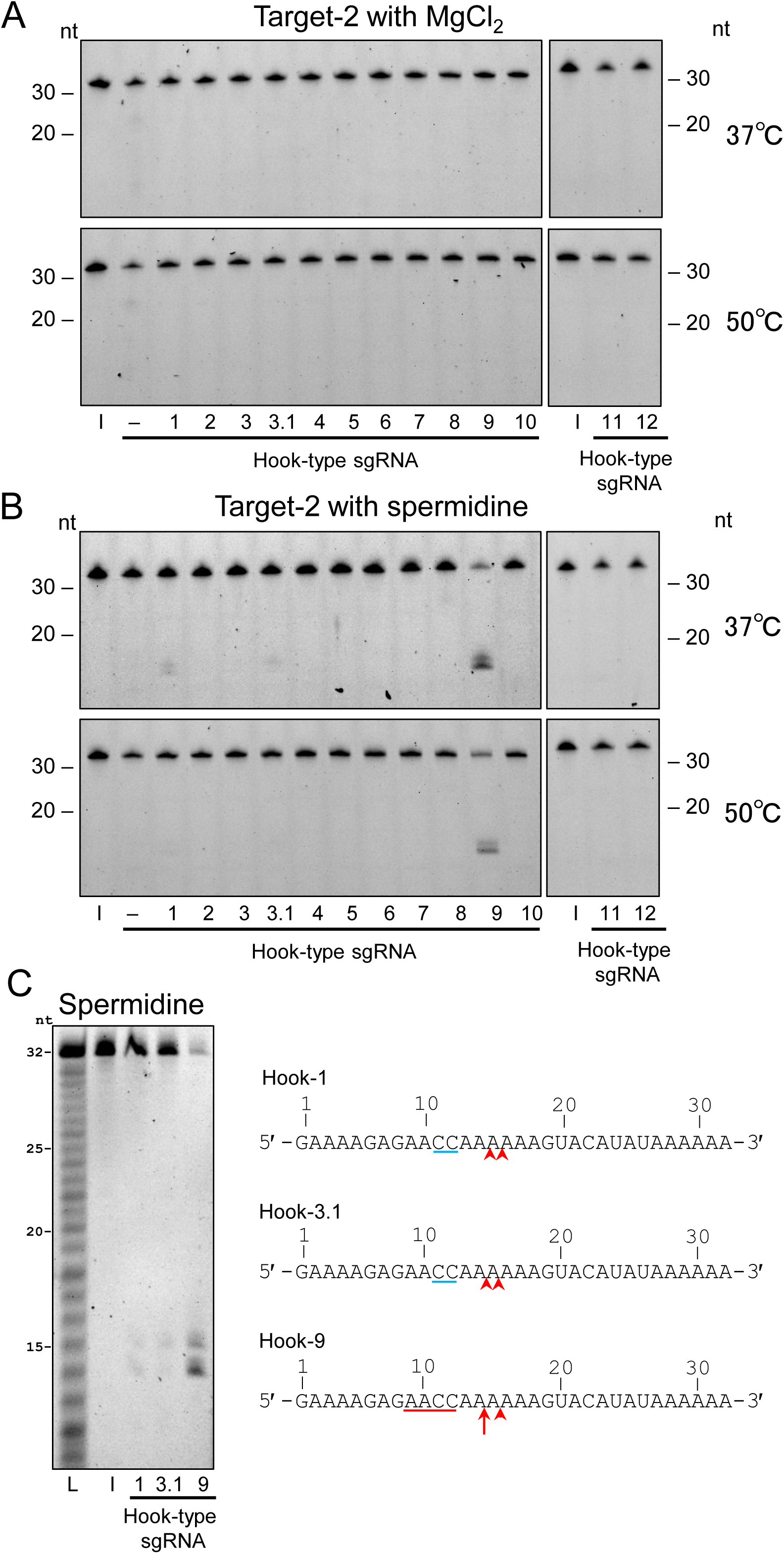
The tRNase Z^L^ cleavage assay for Target-2 in the presence of the hook-type sgRNAs Hook-1–Hook-12. The cleavage assays were performed with 5 mM MgCl_2_. (A) or 0.2 mM spermidine (B) at 37 or 50°C, and the reaction products were separated on a denaturing 12.5% polyacrylamide gel. (C) The cleaved RNA products in the reactions with spermidine at 37°C were also separated on a denaturing 20% polyacrylamide sequencing gel. The sites of cleavages with spermidine are indicated by red arrows/arrowheads on the target RNA. A relatively strong cleavage site is denoted by an arrow. The potential sgRNA binding sequences of ≧ 4 nt and < 4 nt are indicated by red and blue lines, respectively. 20- and 30-nt RNAs and an alkaline ladder were used as size standards. L, alkaline ladder; I, input RNA.

### Cleavage assays for unstructured Target-3 and Target-4

Target-3 had seven potential sgRNA binding sites: one for Hook-3, Hook-3.1, Hook-4, Hook-6 and Hook-10 and two for Hook-5 and Hook-9 (S3–S7 Fig., S10 Fig. and S11 Fig.). This target was cleaved by Hook-4-guided tRNase Z^L^ 1-nt downstream of a D-nt (the 22nd nt G) and by Hook-9-guided tRNase Z^L^ 1/2-nt downstream of a D-nt (the 13th nt A) only in the presence of spermidine (Fig 5, S5 Fig. and S10 Fig.).

**Fig 5.**
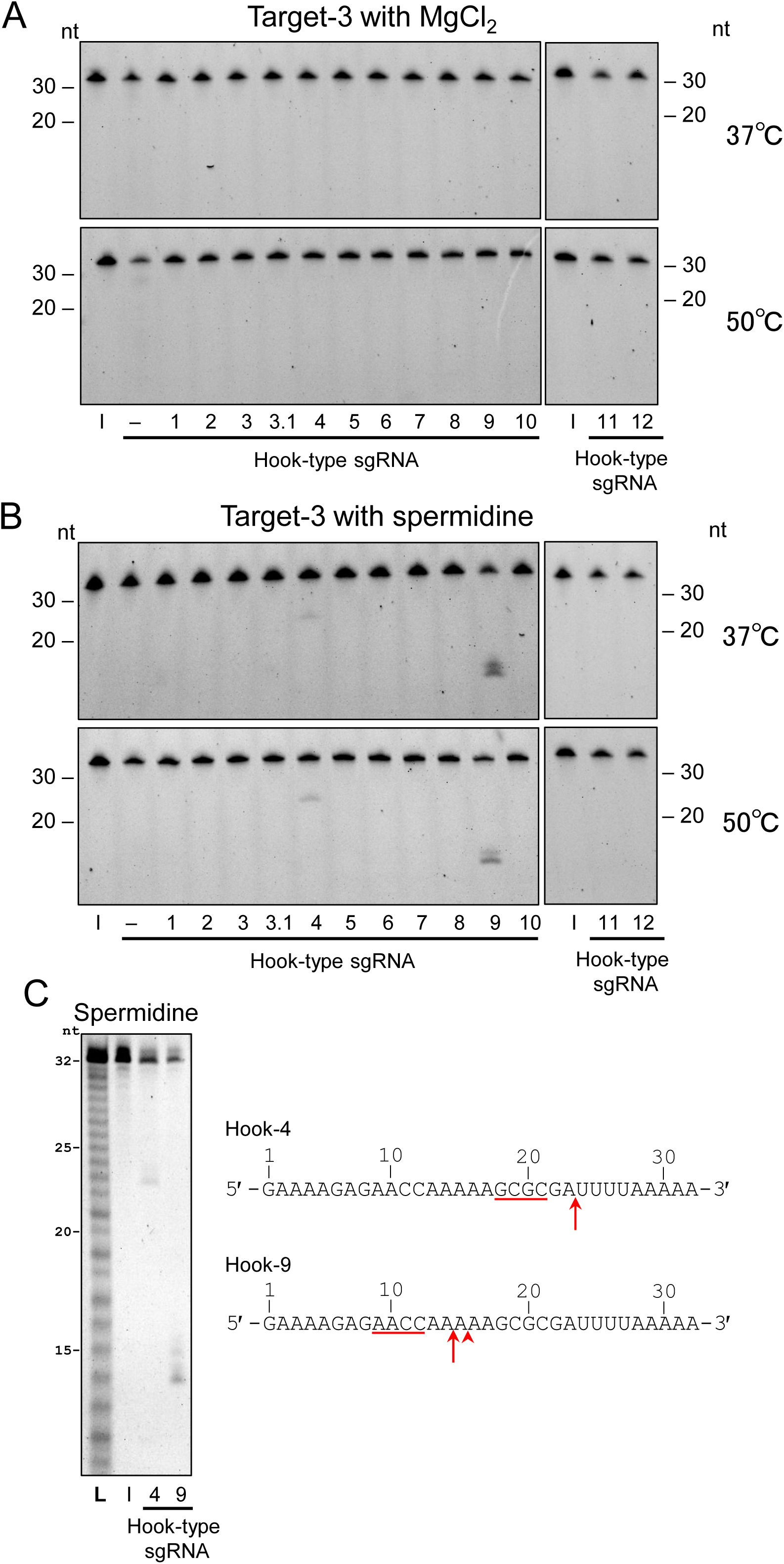
The tRNase Z^L^ cleavage assay for Target-3 in the presence of the hook-type sgRNAs Hook-1–Hook-12. The cleavage assays were performed with 5 mM MgCl_2_. (A) or 0.2 mM spermidine (B) at 37 or 50°C, and the reaction products were separated on a denaturing 12.5% polyacrylamide gel. (C) The cleaved RNA products in the reactions with spermidine at 37°C were also separated on a denaturing 20% polyacrylamide sequencing gel. The sites of cleavages with spermidine are indicated by red arrows/arrowheads on the target RNA. A relatively strong cleavage site is denoted by an arrow. The potential sgRNA binding sequences are indicated by red lines. 20- and 30-nt RNAs and an alkaline ladder were used as size standards. L, alkaline ladder; I, input RNA.

Target-4 had four potential sgRNA binding sites: one for Hook-11 and Hook-12 and three for Hook-10 (S11–S13 Fig.). Target-4 was cleaved only by Hook-11-guided tRNase Z^L^ 1/2-nt downstream of a D-nt (the 12th nt A) (Fig 6 and S12 Fig.).

**Fig 6.**
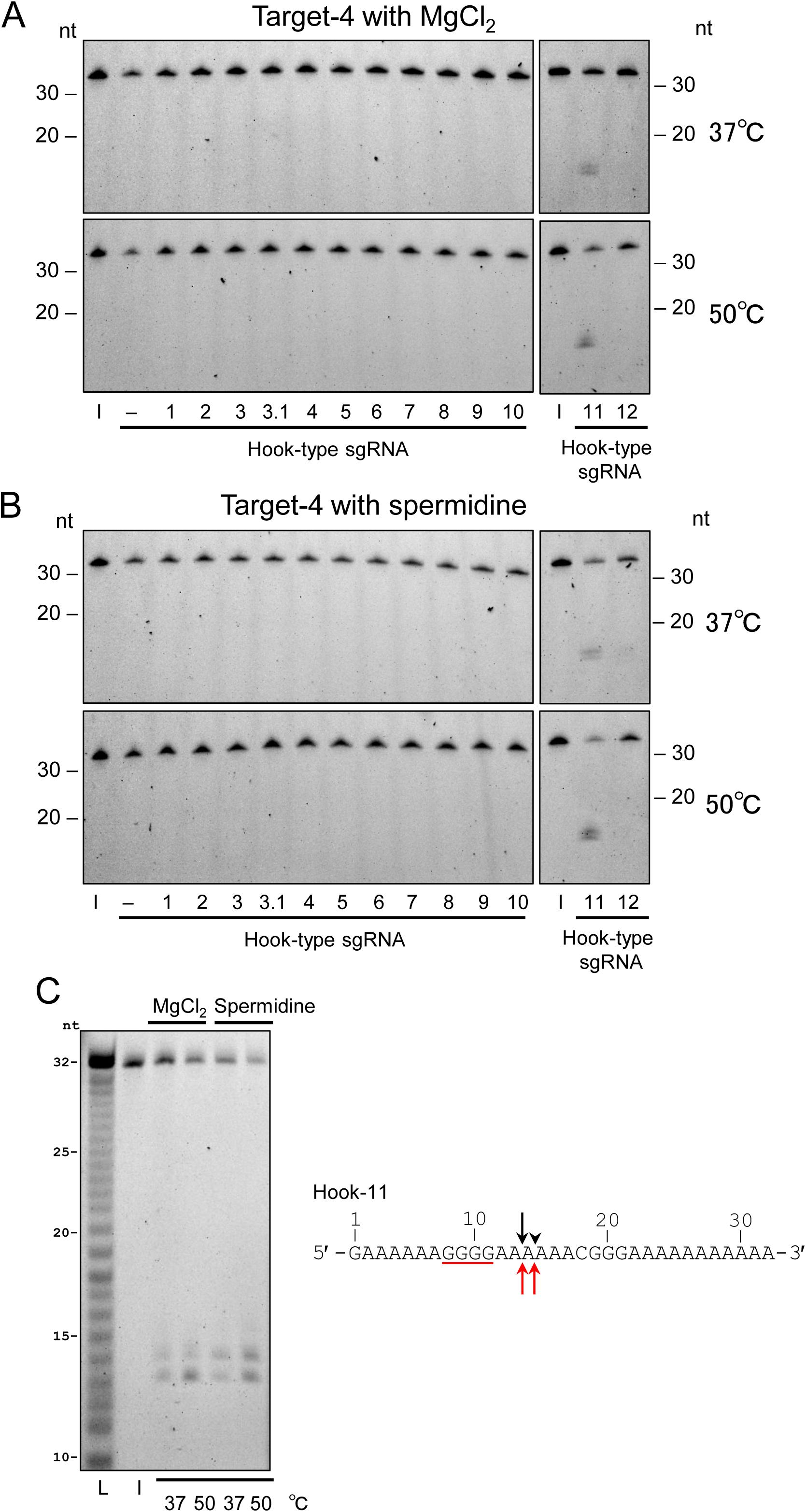
The tRNase Z^L^ cleavage assay for Target-4 in the presence of the hook-type sgRNAs Hook-1–Hook-12. The cleavage assays were performed with 5 mM MgCl_2_. (A) or 0.2 mM spermidine (B) at 37 or 50°C, and the reaction products were separated on a denaturing 12.5% polyacrylamide gel. (C) The cleaved RNA products were also separated on a denaturing 20% polyacrylamide sequencing gel. The sites of cleavages with MgCl_2_ and with spermidine are indicated by black and red arrows/arrowheads, respectively, on the target RNA. A relatively strong cleavage site is denoted by an arrow. The potential sgRNA binding sequence is indicated by a red line. 20- and 30-nt RNAs and an alkaline ladder were used as size standards. L, alkaline ladder; I, input RNA.

### Cleavage assays for tightly-folded Target-5

Target-5 contained nine potential sgRNA binding sites: one for Hook-1, Hook-10, Hook-11 and Hook-12, two for Hook-2 and Hook-3.1, three for Hook-9 and four for Hook-3 (S1–S4 Fig. and S10–S13 Fig.). However, Target-5 was not cleaved by any sgRNA-guided tRNase Z^L^ under any conditions (Fig 7).

**Fig 7.**
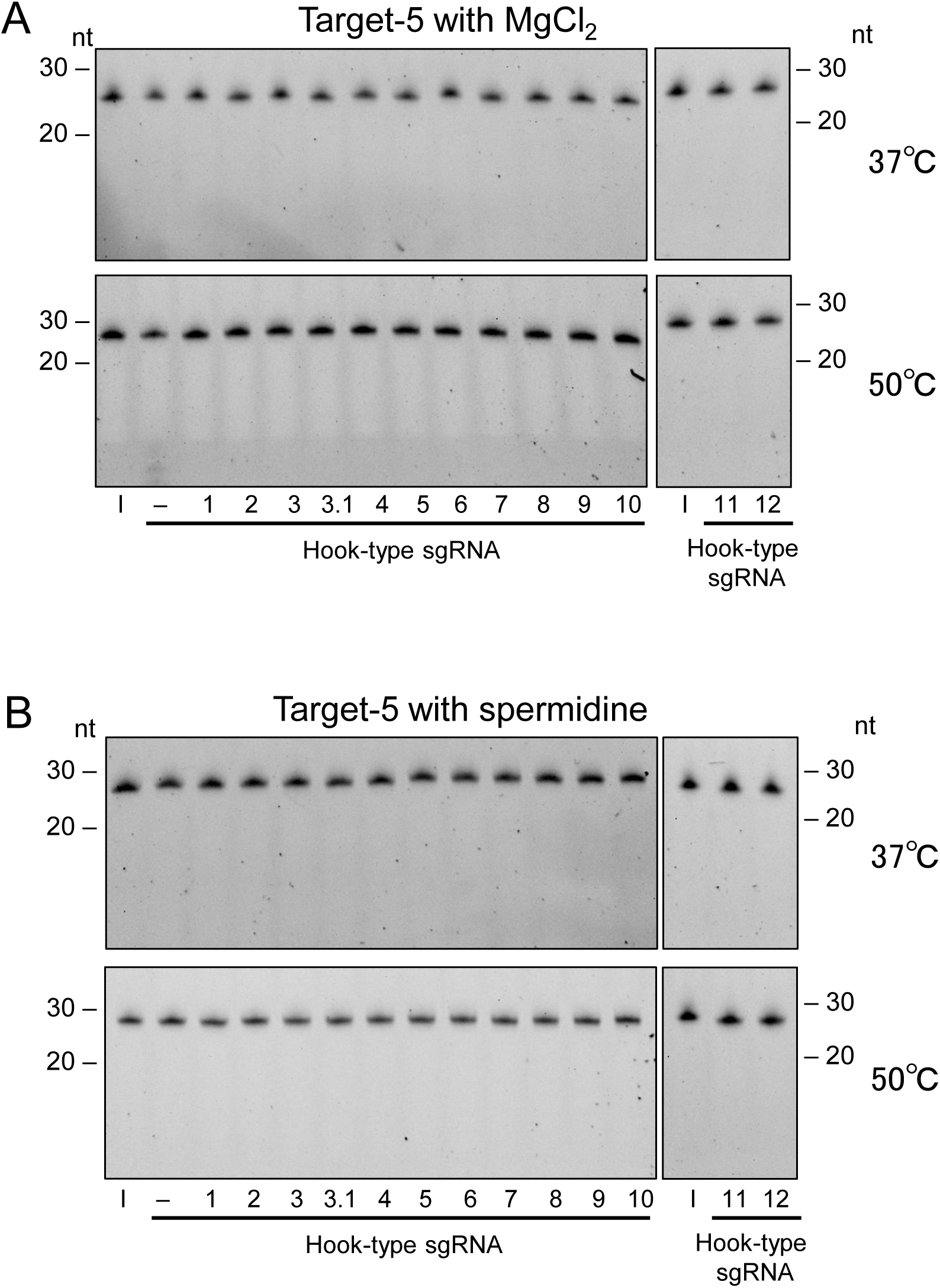
The tRNase Z^L^ cleavage assay for Target-5 in the presence of the hook-type sgRNAs Hook-1–Hook-12. The cleavage assays were performed with 5 mM MgCl_2_. (A) or 0.2 mM spermidine (B) at 37 or 50°C, and the reaction products were separated on a denaturing 20% polyacrylamide gel. 20- and 30-nt RNAs were used as size standards. L, alkaline ladder; I, input RNA.

### Cleavage assays for loosely-folded Target-6–Target-9

Target-6 had nine potential sgRNA binding sites: one for Hook-1, Hook-2, Hook-4 and Hook-10, two for Hook-3.1, three for Hook-3 and four for Hook-9 (S1–S5 Fig., S10 Fig. and S11 Fig.). Target-6 was cleaved by Hook-1-, Hook-2-, and Hook-3-guided tRNase Z^L^ after the 13th nt G (D-nt) and 1-nt downstream of it (Fig 8, S1 Fig., S2 Fig. and S3 Fig.). Hook-3.1-guided tRNase Z^L^ cleaved Target-6 around the 13th nt G, suggesting that the target RNA can be recognized by Hook-3.1-guided tRNase Z^L^ through three consecutive G-C WCF base-pairings (Fig 8 and S4 Fig.). The cleavage after the 21st nt U (D-nt) was also observed in the reaction with spermidine by Hook-3-guided tRNase Z^L^ and in the reaction with MgCl_2_ by Hook-3.1-guided tRNase Z^L^ (Fig 8, S3 Fig., and S4 Fig.).

**Fig 8.**
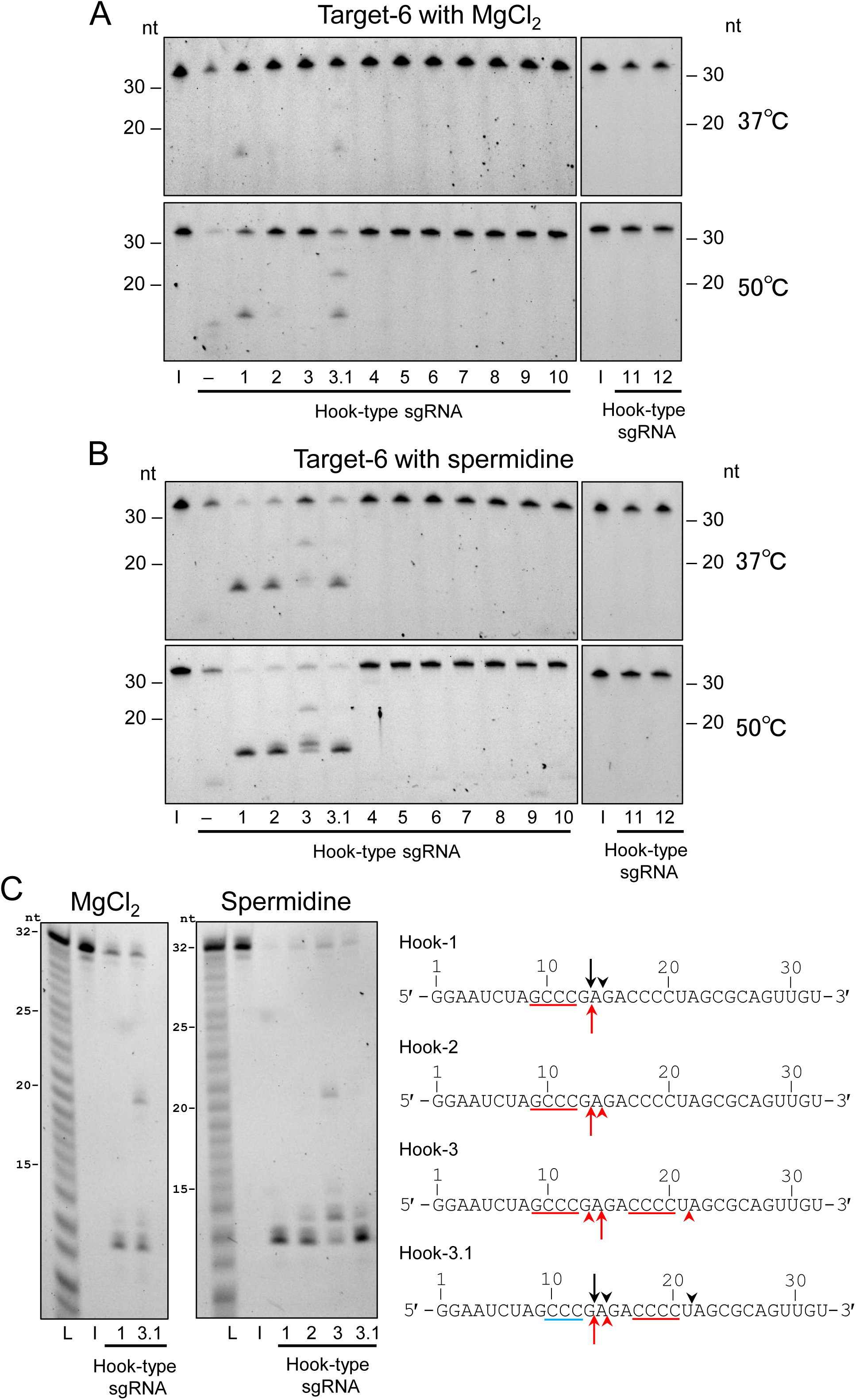
The tRNase Z^L^ cleavage assay for Target-6 in the presence of the hook-type sgRNAs Hook-1–Hook-12. The cleavage assays were performed with 5 mM MgCl_2_. (A) or 0.2 mM spermidine (B) at 37 or 50°C, and the reaction products were separated on a denaturing 20% polyacrylamide gel. (C) The cleaved RNA products in the reactions at 50°C were also separated on a denaturing 12.5% polyacrylamide sequencing gel. The sites of cleavages with MgCl_2_ and with spermidine are indicated by black and red arrows/arrowheads, respectively, on the target RNA. A relatively strong cleavage site is denoted by an arrow. The potential sgRNA binding sequences of ≧ 4 nt and < 4 nt are indicated by red and blue lines, respectively. 20- and 30-nt RNAs and an alkaline ladder were used as size standards. L, alkaline ladder; I, input RNA.

Target-7 had thirteen potential sgRNA binding sites: one for Hook-3.1, Hook-4, and Hook-8, two for Hook-1, Hook-2 and Hook-9, three for Hook-6, Hook-7 and Hook-10 and four for Hook-3 and Hook-5 (S1–S11 Fig.). Target-7 was cleaved only by Hook-11-guided tRNase Z^L^ around the 8th nt A in the presence of spermidine (Fig 9 and S12 Fig.). This suggests that the target RNA can be recognized by Hook-11-guided tRNase Z^L^ via three consecutive G-C WCF base-pairings.

**Fig 9.**
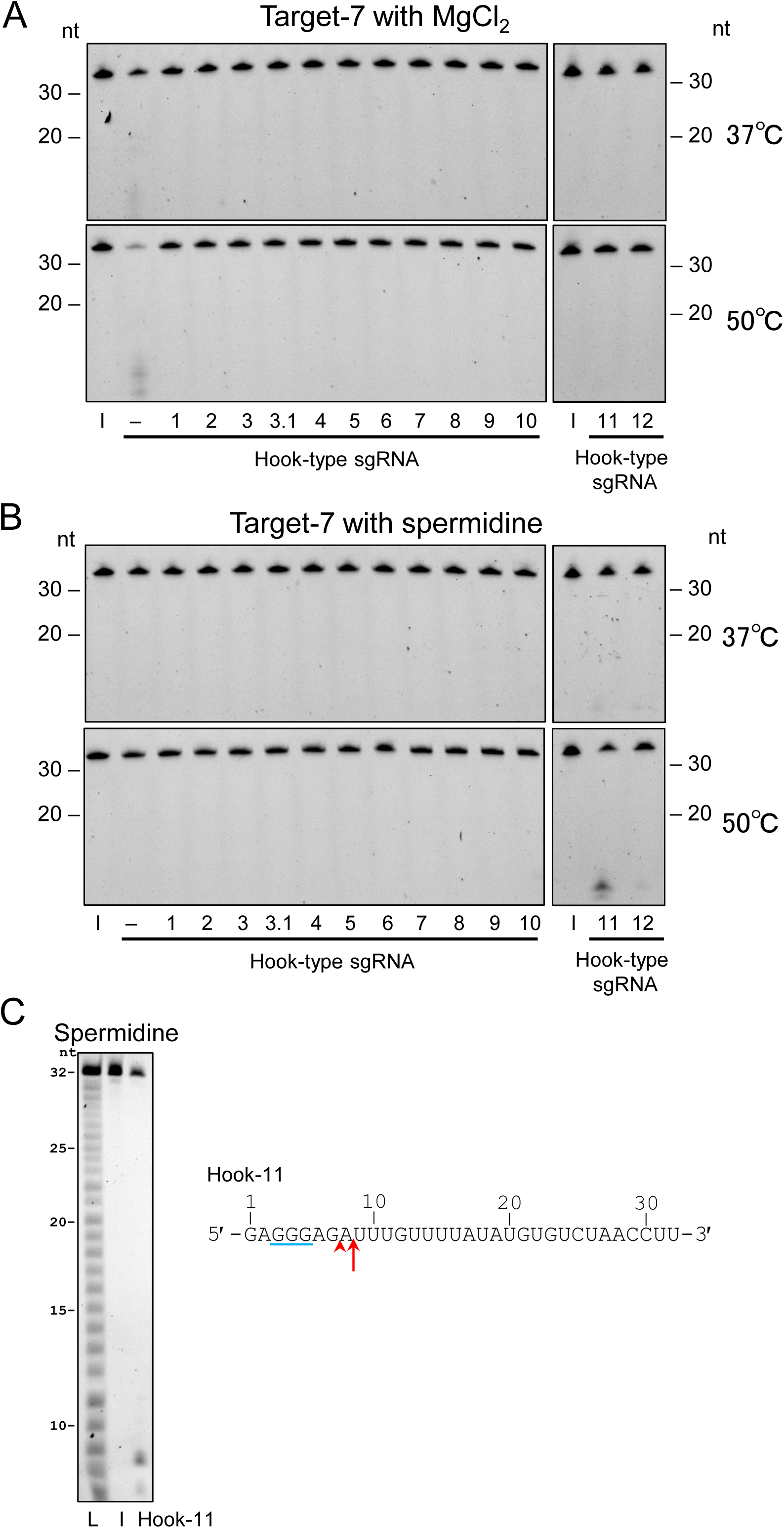
The tRNase Z^L^ cleavage assay for Target-7 in the presence of the hook-type sgRNAs Hook-1–Hook-12. The cleavage assays were performed with 5 mM MgCl_2_. (A) or 0.2 mM spermidine (B) at 37 or 50°C, and the reaction products were separated on a denaturing 20% polyacrylamide gel. (C) The cleaved RNA products in the reactions at 50°C were also separated on a denaturing 12.5% polyacrylamide sequencing gel. The sites of cleavages with spermidine are indicated by a red arrow/arrowhead on the target RNA. A relatively strong cleavage site is denoted by an arrow. The potential sgRNA binding sequence is indicated by a blue line. 20- and 30-nt RNAs and an alkaline ladder were used as size standards. L, alkaline ladder; I, input RNA.

Target-8 had nine potential sgRNA binding sites: one for Hook-4, Hook-11 and Hook-12, two for Hook-2, Hook-5, Hook-8 and Hook-10, and four for Hook-9 (S2 Fig., S5 Fig., S6 Fig., S9–S13 Fig.). Target-8 was cleaved only by Hook-12-guided tRNase Z^L^ 1-nt downstream of a D-nt (the 13th nt G) in the presence of spermidine (Fig 10 and S13 Fig.).

**Fig 10.**
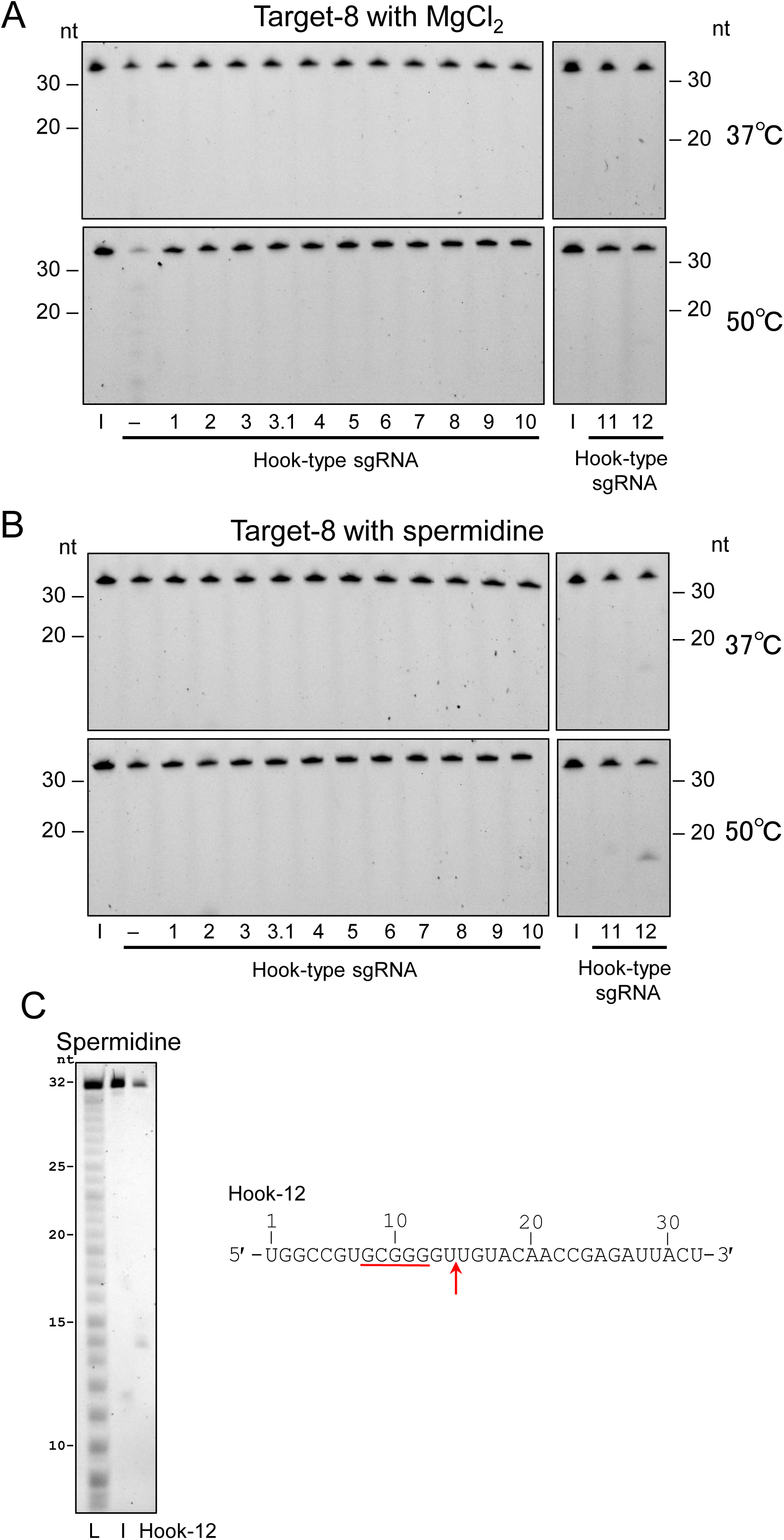
The tRNase Z^L^ cleavage assay for Target-8 in the presence of the hook-type sgRNAs Hook-1–Hook-12. The cleavage assays were performed with 5 mM MgCl_2_. (A) or 0.2 mM spermidine (B) at 37 or 50°C, and the reaction products were separated on a denaturing 20% polyacrylamide gel. (C) The cleaved RNA products in the reactions at 50°C were also separated on a denaturing 12.5% polyacrylamide sequencing gel. The site of cleavage with spermidine is indicated by a red arrow on the target RNA. The potential sgRNA binding sequence is indicated by a red line. 20- and 30-nt RNAs and an alkaline ladder were used as size standards. L, alkaline ladder; I, input RNA.

Target-9 had fifteen potential sgRNA binding sites: one for Hook-10, two for Hook-1 and Hook-3.1, three for Hook-4, five for Hook-3 and six for Hook-2 (S1–S5 Fig., and S11 Fig.). Hook-1- and Hook-2-guided tRNase Z^L^ cleaved Target-9 3-nt downstream of a D-nt (the 13th nt G) under both MgCl_2_ and spermidine conditions and after the 20th nt C (D-nt) only under the spermidine conditions (Fig 11, S1 Fig. and S2 Fig.). Hook-3 barely guided tRNase Z^L^ to cleave Target-9 3-nt downstream of a D-nt (the 13th nt G) and after the 21st nt U (D-nt), and Hook-3.1 also barely guided tRNase Z^L^ to cleave the target after the 21st nt U (D-nt) and after the 16th nt G (Fig 11, S3 Fig. and S4 Fig.). Target-9 was also cleaved faintly by Hook-4-guided tRNase Z^L^ after the 27th nt A (D-nt) (Fig 11 and S5 Fig.).

**Fig 11.**
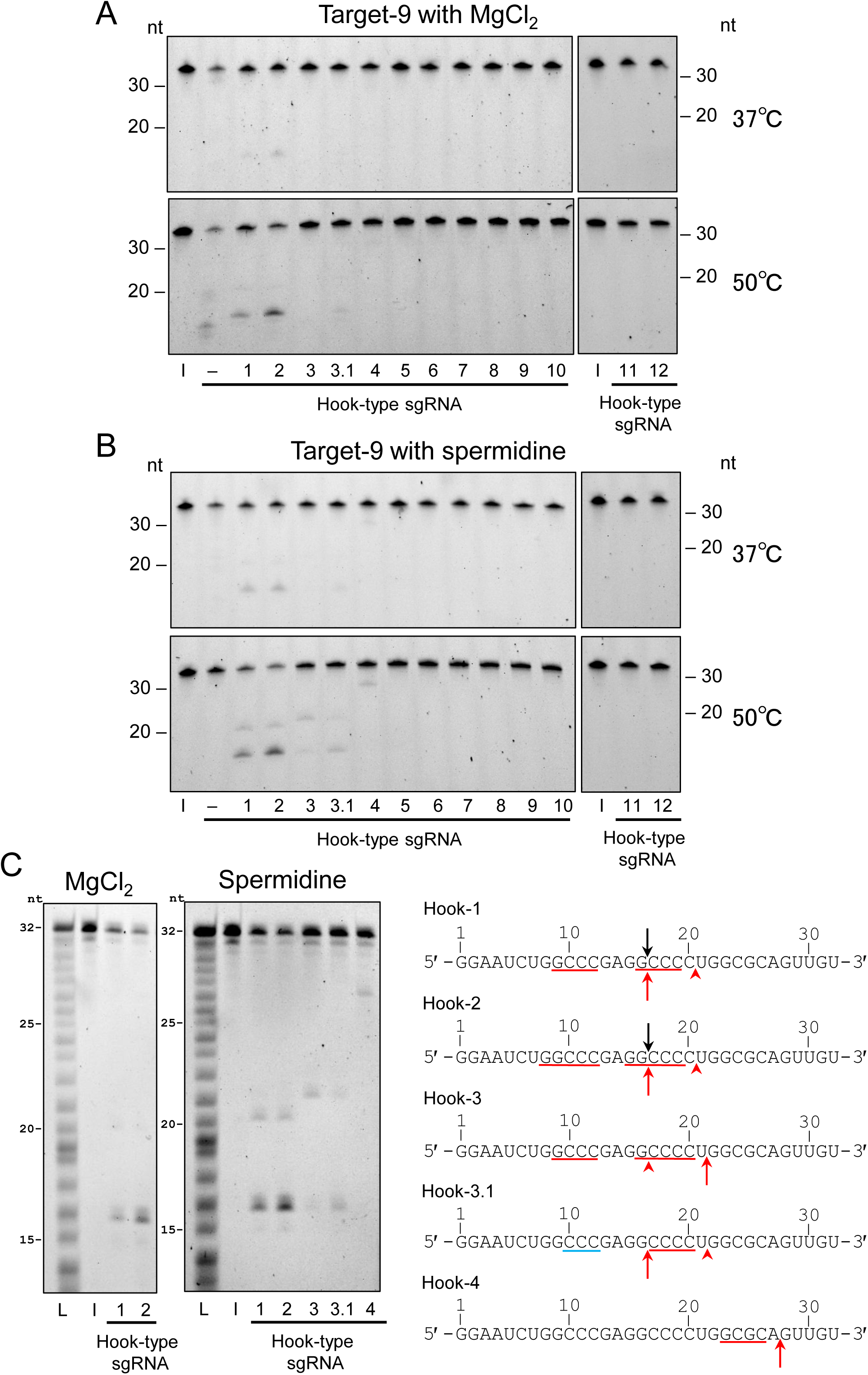
The tRNase Z^L^ cleavage assay for Target-9 in the presence of the hook-type sgRNAs Hook-1–Hook-12. The cleavage assays were performed with 5 mM MgCl_2_. (A) or 0.2 mM spermidine (B) at 37 or 50°C, and the reaction products were separated on a denaturing 20% polyacrylamide gel. (C) The cleaved RNA products in the reactions at 50°C were also separated on a denaturing 12.5% polyacrylamide sequencing gel. The sites of cleavages with MgCl_2_ and with spermidine are indicated by black and red arrows/arrowheads, respectively, on the target RNA. A relatively strong cleavage site is denoted by an arrow. The potential sgRNA binding sequences of ≧ 4 nt and < 4 nt are indicated by red and blue lines, respectively. 20- and 30-nt RNAs and an alkaline ladder were used as size standards. L, alkaline ladder; I, input RNA.

## Discussion

In this study, we examined nine 32-nt RNA targets of various degrees of folding for cleavability by human tRNase Z^L^ guided by each of thirteen hook-type sgRNAs. The assays were carried out under four different conditions: at 37 or 50°C with MgCl_2_ or spermidine. And the cleavage sites were determined by analyzing the cleavage products on a sequencing gel. On the whole, the cleavages of target RNA were more efficient at 50°C than at 37°C and with spermidine than with MgCl_2_. From these 468 assays, we identified four valuable findings as described below.

Hook-5–Hook-10, which contain two to four A or U nt in the target RNA recognition sequence (Fig 2), were not able to guide tRNase Z^L^ to cleave any of the nine RNA targets including unstructured Target-1–Target-4 with the exception of Hook-9 (S6–S11 Fig.). These observations suggest that at least two consecutive G or C nt in the target RNA recognition sequence are needed for hook-type sgRNA to work as a guide for tRNase Z^L^ (Fig 2).

Hook-1 and Hook-2 that has a one-base-pair shorter stem structure than Hook-1 contain the common target RNA recognition sequence 5′-GGGC-3′, and Hook-3.1 and Hook-3 that has a one-base-pair shorter stem structure than Hook-3.1 contain the common target RNA recognition sequence 5′-GGGG-3′ (Fig 2). These sgRNAs were designed to examine how the one-base-pair difference affects their guiding efficiency. The assays with Target-1 and Target-6, both of which contained their target sequences 5′-GCCC-3′ and 5′-CCCC-3′, showed that, on the whole, Hook-1 and Hook-3.1 were more efficient in the guiding ability than Hook-2 and Hook-3, respectively, especially under the conditions with MgCl_2_ (Fig 3 and Fig 8).

Hook-1 and Hook-3.1, but not Hook-2 and Hook-3, also barely guided the cleavage of Target-2 that contained neither 5′-GCCC-3′ nor 5′-CCCC-3′, probably through recognizing two consecutive C nt at the 11th and 12th positions (Fig 4). Similarly, Hook-1 guided the cleavage of Target-1 around the 12th nt A much more efficiently than Hook-2, probably through recognizing three consecutive C nt at the 8th– 10th or 9th–11th positions (Fig 3).

Target-9 contained the five nt sequences 5′-GGCCC-3′ (at the 8th–12th and 15th–19th positions) and 5′-GCCCC-3′ (at the 16th–20th positions) that are complementary to the five nt 5′-end single-stranded regions of Hook-2 and Hook-3, respectively (Fig 11). Hook-2 guided the cleavage 4-nt downstream of 5′-GGCCC-3′ (at the 8th–12th positions) better than Hook-1, and the guiding efficiency at the other site (the 15th–19th positions) was not different between the two sgRNAs (Fig 11). Similarly, Hook-3 guided the cleavage downstream of 5′-GCCCC-3′ (at the 16th–20th positions) better than Hook-3.1.

Taken together, these observations with respect to the assays for Hook-1 vs Hook-2 and Hook-3.1 vs Hook-3 would lead to the following two rules (the second finding): A longer target RNA recognition sequence of hook-type sgRNA provides better guiding ability. Hook-type sgRNA that can form with target RNA a hairpin structure without bulged nucleotides shows a better guiding ability. There was one exception to the first rule: Hook-3 guided the cleavages after 5′-GCCC-3′ (at the 9th– 12th positions) in Target-6 and Target-9 less efficiently than Hook-3.1 (Fig 8 and Fig 11). This may be attributed to a less stable interaction between tRNase Z^L^ and the RNA complex of Hook-3 with Target-6 or Target-9.

The third finding is that tRNase Z^L^ with hook-type sgRNA, which was designed to work as a 4-base-recognizing RNA cutter, can also work efficiently as a 3-base-recognizing RNA cutter in some cases (Fig 3, Fig 8 and Fig 9).

The last and most important finding is that cleavage of target RNA by tRNase Z^L^ guided by hook-type sgRNA occurs only in unstructured or loosely-folded target RNA (Fig 1). From this finding and the previous observation [16], it can be deduced that sgRNA-guided tRNase Z^L^ can discriminate less folded regions from tightly folded ones in long cellular mRNA or non-coding RNA.

Structure probing of cellular RNA by introducing into the cells the hook-type sgRNA designed based on the findings of this study and subsequently analyzing the resulting RNA fragments by a next-generation sequencer would provide valuable information on the higher-order structures of the cellular RNA. And this information would reinforce the data obtained by other methods such as chemical probing. In addition, the structure probing with sgRNA-guided tRNase Z^L^ would directly provide good target sites not only for TRUE gene silencing but also for gene suppression with conventional ASO and RNA interference, because this method provides information on accessibility of relatively large nucleic-acids/protein-enzyme complexes such sgRNA/ tRNase Z^L^, ASO/RNase HI, and siRNA/AGO2, which cannot be provided directly by chemical probing. We are currently planning to carry out *in vivo* structure probing of the MALAT1 RNA by introducing several hook-type sgRNAs into A549 cells to find better target sites for TRUE gene silencing.

## Author Contributions

**Conceptualization:** Masayuki Nashimoto.

**Data curation:** Masayuki Takahashi, Masayuki Nashimoto.

**Formal analysis:** Masayuki Takahashi, Masayuki Nashimoto.

**Funding acquisition:** Masayuki Nashimoto.

**Investigation:** Masayuki Takahashi.

**Project administration:** Masayuki Nashimoto.

**Resources:** Masayuki Takahashi, Masayuki Nashimoto.

**Supervision:** Masayuki Nashimoto.

**Validation:** Masayuki Takahashi, Masayuki Nashimoto.

**Visualization:** Masayuki Takahashi.

**Writing – original draft:** Masayuki Nashimoto.

**Writing – review & editing:** Masayuki Takahashi, Masayuki Nashimoto.

## Data Availability Statement

All relevant data are within the manuscript.

## Supporting Information

**S1 Fig. Predicted secondary structures of RNA targets, Target-1–Target-9, with the potential Hook-1 binding sites for them.** The potential Hook-1 binding sequences are indicated by red lines. The binding is thought to occur through base pairs including the non-WCF G-U base pair. The target RNA recognition sequence of Hook-1 is 5′-GGGC-3′. An arrow indicates the position where cleavage by tRNase Z^L^ occurred under at least one of the four assay conditions.

**S2 Fig. Predicted secondary structures of RNA targets, Target-1–Target-9, with the potential Hook-2 binding sites for them.** The potential Hook-2 binding sequences are indicated by red lines. The binding is thought to occur through base pairs including the non-WCF G-U base pair. The target RNA recognition sequence of Hook-2 is 5′-GGGC-3′, 5′-GGCC-3′ or 5′-GGGCC-3′. An arrow indicates the position where cleavage by tRNase Z^L^ occurred under at least one of the four assay conditions.

**S3 Fig. Predicted secondary structures of RNA targets, Target-1–Target-9, with the potential Hook-3 binding sites for them.** The potential Hook-3 binding sequences are indicated by red lines. The binding is thought to occur through base pairs including the non-WCF G-U base pair. The target RNA recognition sequence of Hook-3 is 5′-GGGG-3′, 5′-GGGC-3′ or 5′-GGGGC-3′. An arrow indicates the position where cleavage by tRNase Z^L^ occurred under at least one of the four assay conditions.

**S4 Fig. Predicted secondary structures of RNA targets, Target-1–Target-9, with the potential Hook-3.1 binding sites for them.** The potential Hook-3.1 binding sequences are indicated by red lines. The binding is thought to occur through base pairs including the non-WCF G-U base pair. The target RNA recognition sequence of Hook-3.1 is 5′-GGGG-3′. An arrow indicates the position where cleavage by tRNase Z^L^ occurred under at least one of the four assay conditions.

**S5 Fig. Predicted secondary structures of RNA targets, Target-1–Target-9, with the potential Hook-4 binding sites for them.** The potential Hook-4 binding sequences are indicated by red lines. The binding is thought to occur through base pairs including the non-WCF G-U base pair. The target RNA recognition sequence of Hook-4 is 5′-GCGC-3′, 5′-CGCC-3′ or 5′-GCGCC-3′. An arrow indicates the position where cleavage by tRNase Z^L^ occurred under at least one of the four assay conditions.

**S6 Fig. Predicted secondary structures of RNA targets, Target-1–Target-9, with the potential Hook-5 binding sites for them.** The potential Hook-5 binding sequences are indicated by red lines. The binding is thought to occur through base pairs including the non-WCF G-U base pair. The target RNA recognition sequence of Hook-5 is 5′-AAAU-3′, 5′-AAUC-3′ or 5′-AAAUC-3′.

**S7 Fig. Predicted secondary structures of RNA targets, Target-1–Target-9, with the potential Hook-6 binding sites for them.** The potential Hook-6 binding sequences are indicated by red lines. The target RNA recognition sequence of Hook-6 is 5′-AAAA-3′, 5′-AAAC-3′ or 5′-AAAAC-3′.

**S8 Fig. Predicted secondary structures of RNA targets, Target-1–Target-9, with the potential Hook-7 binding sites for them.** The potential Hook-7 binding sequences are indicated by red lines. The binding is thought to occur through base pairs including the non-WCF G-U base pair. The target RNA recognition sequence of Hook-7 is 5′-AUAU-3′.

**S9 Fig. Predicted secondary structures of RNA targets, Target-1–Target-9, with the potential Hook-8 binding sites for them.** The potential Hook-8 binding sequences are indicated by red lines. The binding is thought to occur through base pairs including the non-WCF G-U base pair. The target RNA recognition sequence of Hook-8 is 5′-GUAC-3′.

**S10 Fig. Predicted secondary structures of RNA targets, Target-1–Target-9, with the potential Hook-9 binding sites for them.** The potential Hook-9 binding sequences are indicated by red lines. The binding is thought to occur through base pairs including the non-WCF G-U base pair. The target RNA recognition sequence of Hook-9 is 5′-GGUU-3′. An arrow indicates the position where cleavage by tRNase Z^L^ occurred under at least one of the four assay conditions.

**S11 Fig. Predicted secondary structures of RNA targets, Target-1–Target-9, with the potential Hook-10 binding sites for them. The potential Hook-10 binding** sequences are indicated by red lines. The binding is thought to occur through base pairs including the non-WCF G-U base pair. The target RNA recognition sequence of Hook-10 is 5′-UCUC-3′.

**S12 Fig. Predicted secondary structures of RNA targets, Target-1–Target-9, with the potential Hook-11 binding sites for them.** The potential Hook-11 binding sequences are indicated by red lines. The target RNA recognition sequence of Hook-11 is 5′-CCCC-3′. An arrow indicates the position where cleavage by tRNase Z^L^ occurred under at least one of the four assay conditions.

**S13 Fig. Predicted secondary structures of RNA targets, Target-1–Target-9, with the potential Hook-12 binding sites for them.** The potential Hook-12 binding sequences are indicated by red lines. The binding is thought to occur through base pairs including the non-WCF G-U base pair. The target RNA recognition sequence of Hook-12 is 5′-CCCG-3′, 5′-CCGC-3′ or 5′-CCCGC-3′. An arrow indicates the position where cleavage by tRNase Z^L^ occurred under at least one of the four assay conditions.

**S1_raw_images. Original images for gels.**

