## Supplementary Figures for "RNA structure probing with 4-base-recognizing RNA cutters"

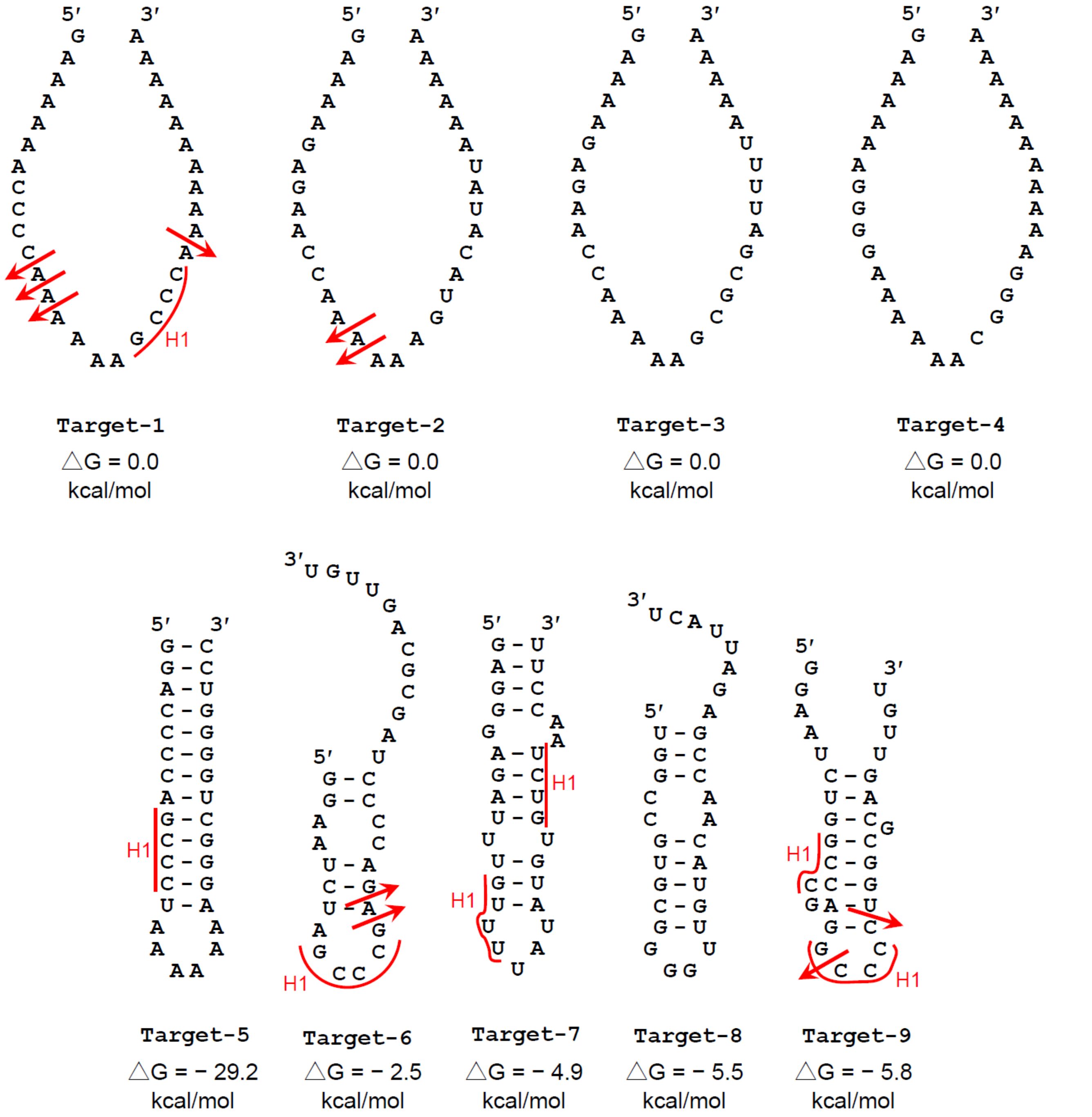
Target-1
△G = 0.0 kcal/mol
Target-2
△G = 0.0 kcal/mol
Target-3
△G = 0.0 kcal/mol
Target-4
△G = 0.0 kcal/mol
Target-5
△G = − 29.2 kcal/mol
Target-6
△G = − 2.5 kcal/mol
Target-7
△G = − 4.9 kcal/mol
Target-8
△G = − 5.5 kcal/mol
Target-9
△G = − 5.8 kcal/mol
H1

S1 Fig.

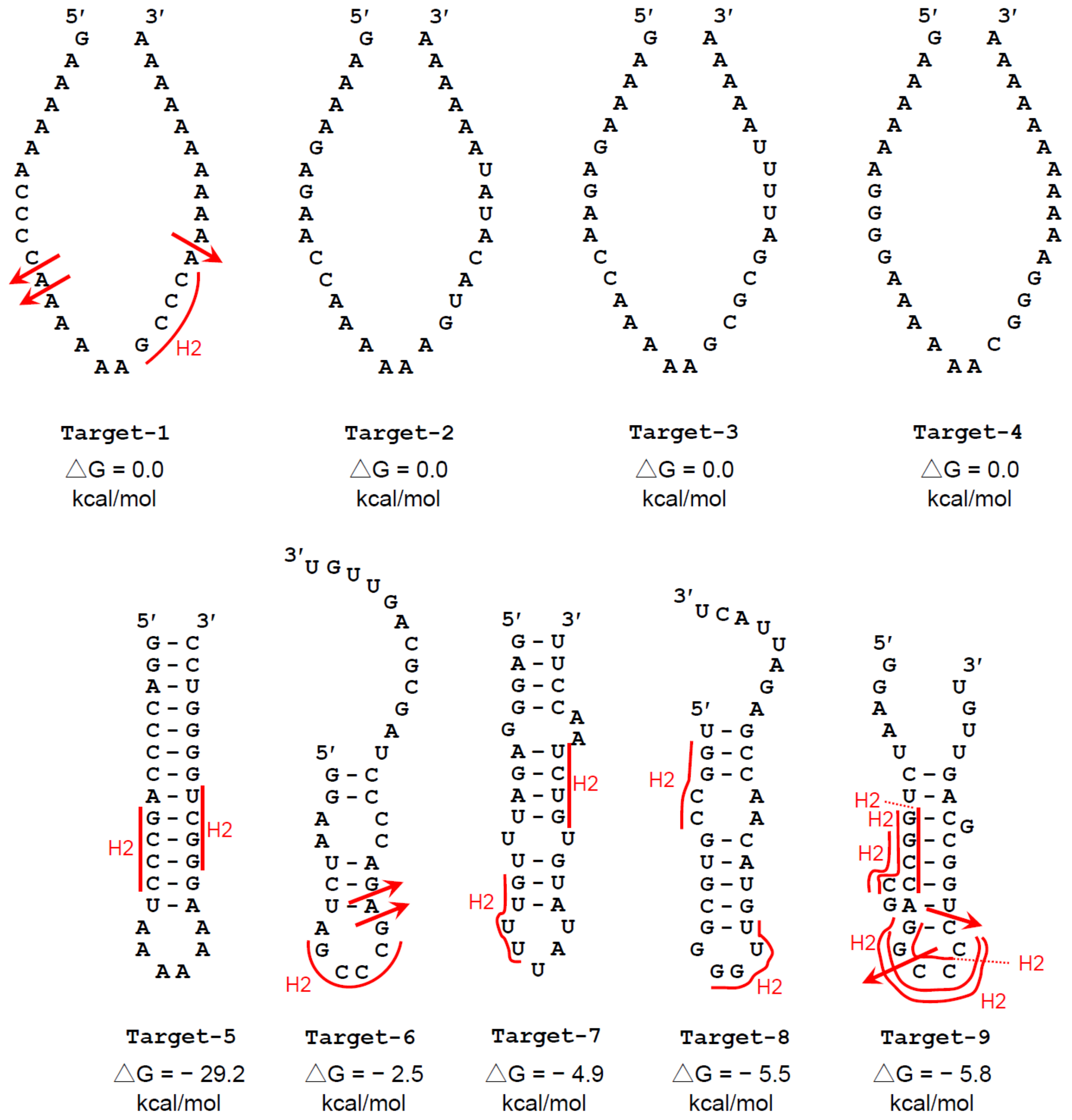

Target-1
△G = 0.0 kcal/mol
Target-2
△G = 0.0 kcal/mol
Target-3
△G = 0.0 kcal/mol
Target-4
△G = 0.0 kcal/mol
Target-5
△G = − 29.2 kcal/mol
Target-6
△G = − 2.5 kcal/mol
Target-7
△G = − 4.9 kcal/mol
Target-8
△G = − 5.5 kcal/mol
Target-9
△G = − 5.8 kcal/mol
H2


S2 Fig.

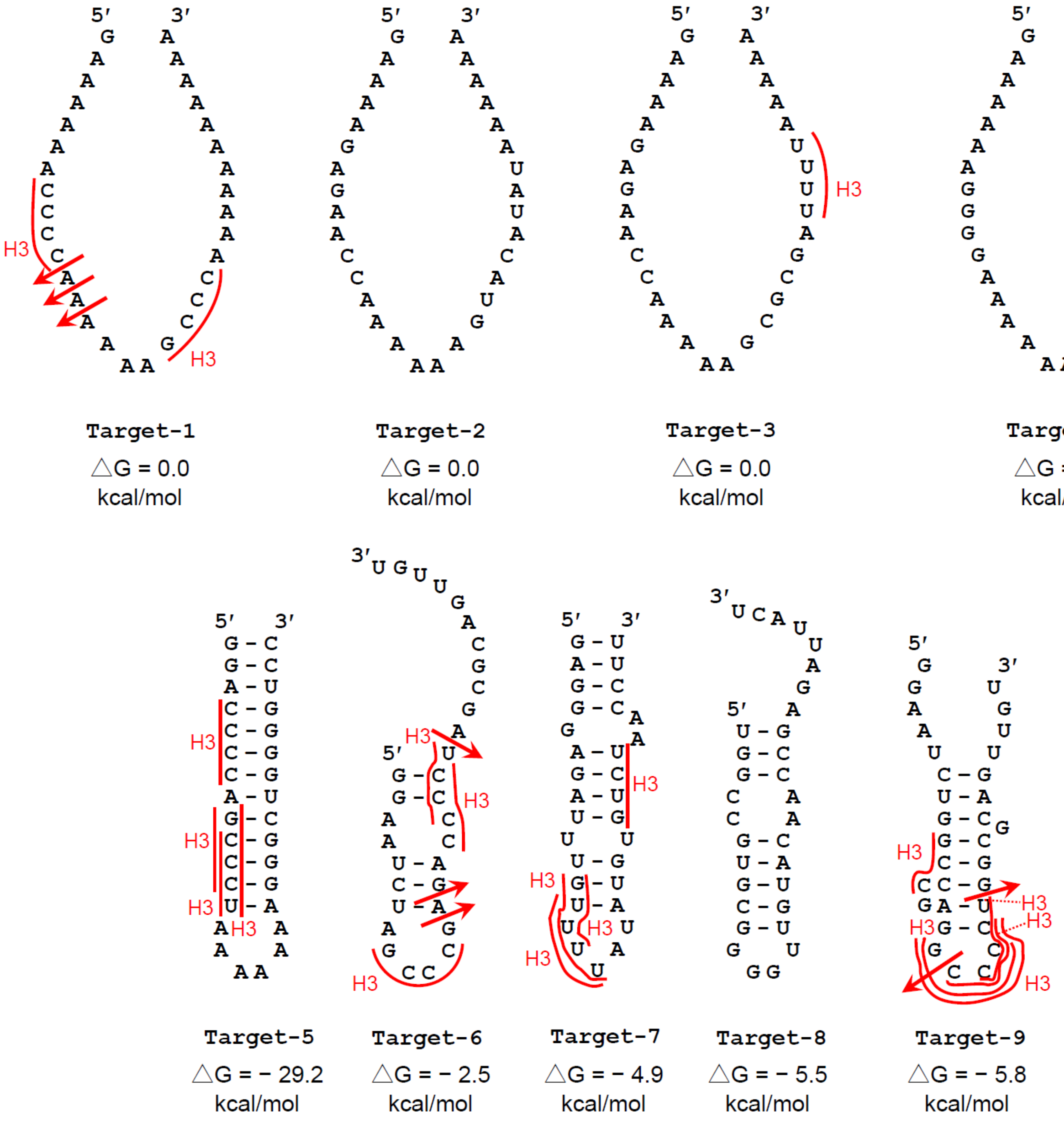
Target-1
△G = 0.0 kcal/mol
Target-2
△G = 0.0 kcal/mol
Target-3
△G = 0.0 kcal/mol
Target-4
△G = 0.0 kcal/mol
Target-5
△G = − 29.2 kcal/mol
Target-6
△G = − 2.5 kcal/mol
Target-7
△G = − 4.9 kcal/mol
Target-8
△G = − 5.5 kcal/mol
Target-9
△G = − 5.8 kcal/mol

S3 Fig.

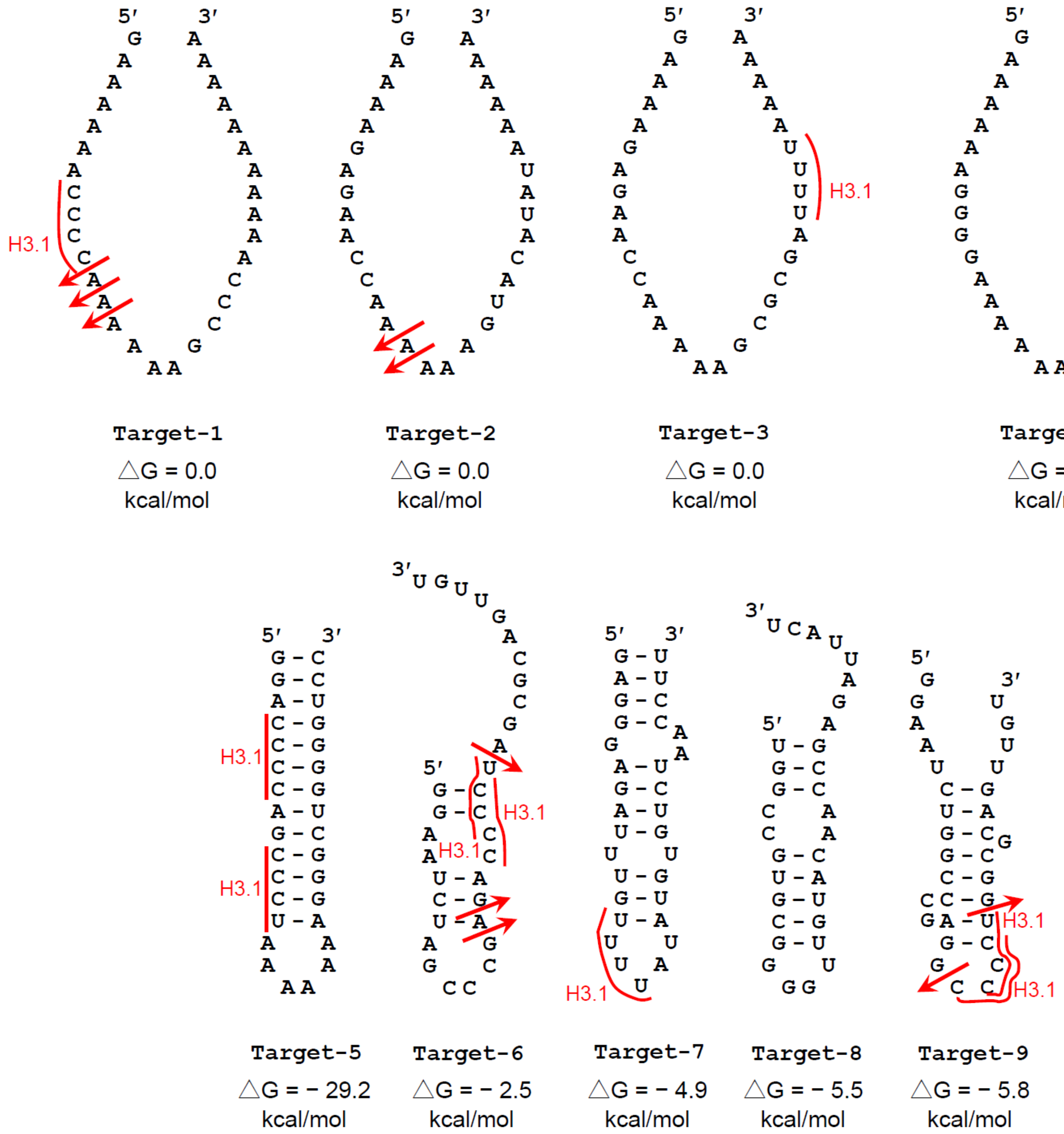

H3.1
Target-1
△G = 0.0 kcal/mol
Target-2
△G = 0.0 kcal/mol
Target-3
△G = 0.0 kcal/mol
Target-4
△G = 0.0 kcal/mol
Target-5
△G = − 29.2 kcal/mol
Target-6
△G = − 2.5 kcal/mol
Target-7
△G = − 4.9 kcal/mol
Target-8
△G = − 5.5 kcal/mol
Target-9
△G = − 5.8 kcal/mol


S4 Fig.

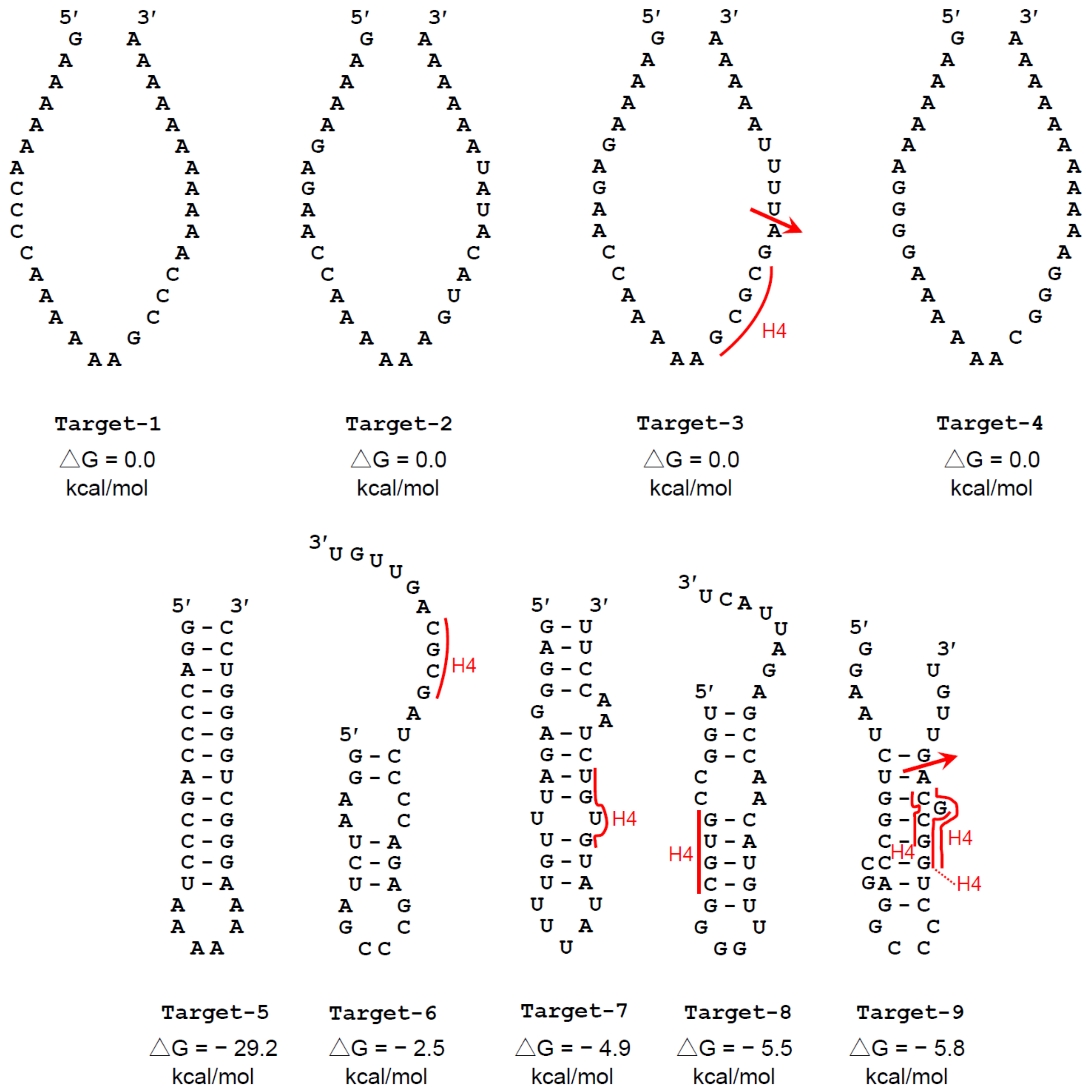
H4
Target-1
△G = 0.0 kcal/mol
Target-2
△G = 0.0 kcal/mol
Target-3
△G = 0.0 kcal/mol
Target-4
△G = 0.0 kcal/mol
Target-5
△G = − 29.2 kcal/mol
Target-6
△G = − 2.5 kcal/mol
Target-7
△G = − 4.9 kcal/mol
Target-8
△G = − 5.5 kcal/mol
Target-9
△G = − 5.8 kcal/mol

S5 Fig.

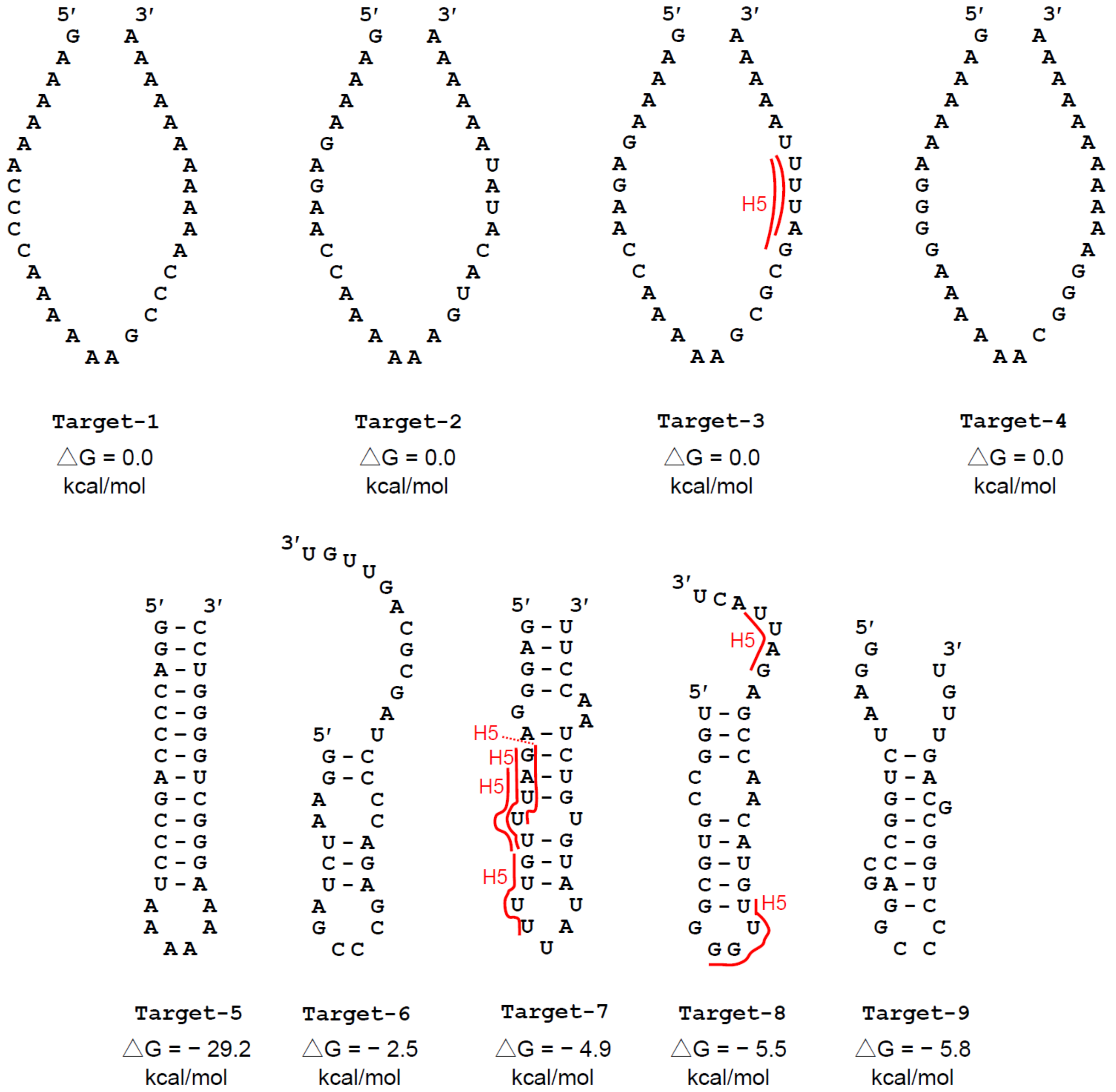

Target-1
△G = 0.0 kcal/mol
Target-2
△G = 0.0 kcal/mol
Target-3
H5
△G = 0.0 kcal/mol
Target-4
△G = 0.0 kcal/mol
Target-5
△G = − 29.2 kcal/mol
Target-6
△G = − 2.5 kcal/mol
Target-7
H5
△G = − 4.9 kcal/mol
Target-8
H5
△G = − 5.5 kcal/mol
Target-9
△G = − 5.8 kcal/mol


S6 Fig.

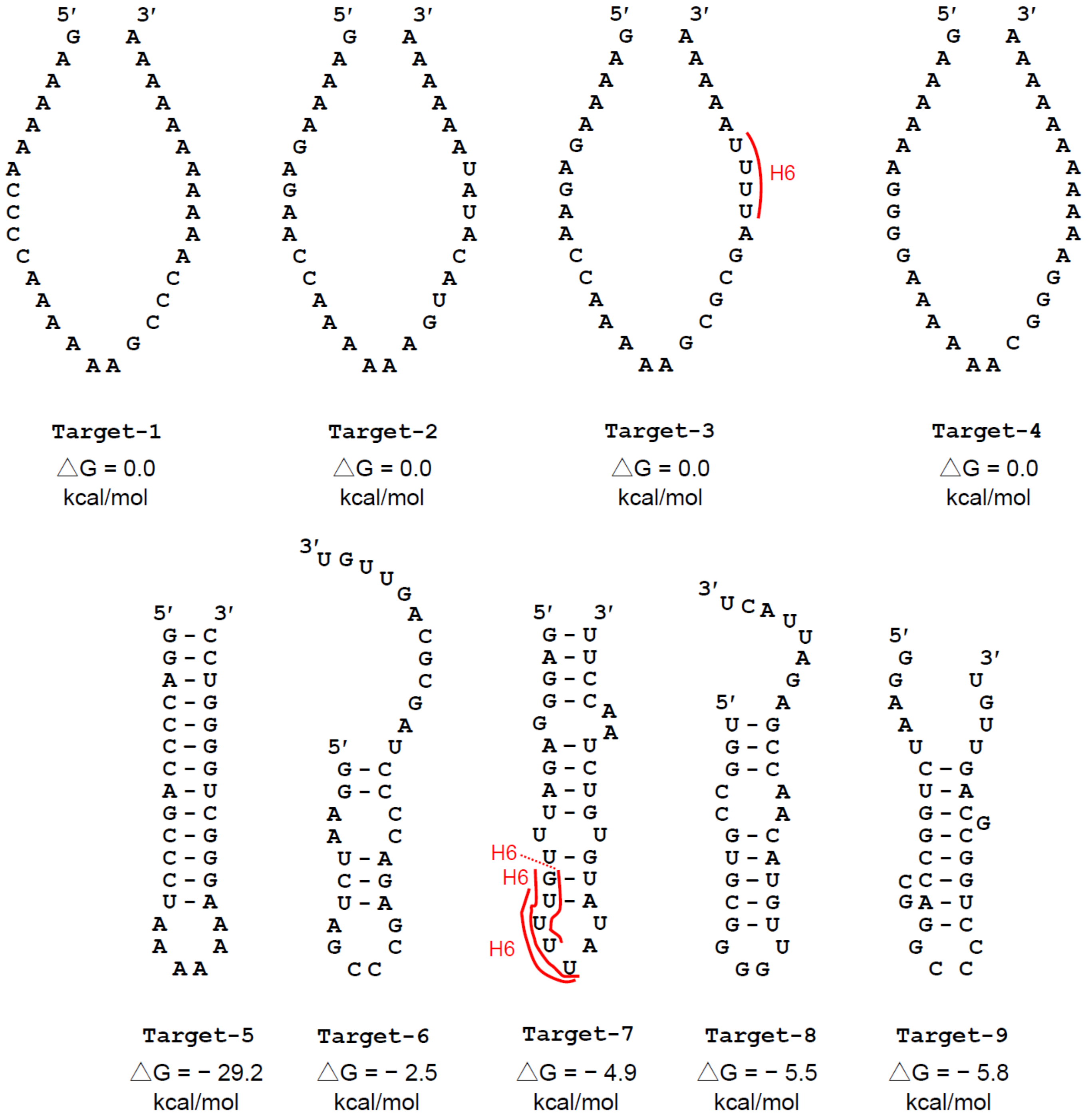

H6
Target-1
△G = 0.0 kcal/mol
Target-2
△G = 0.0 kcal/mol
Target-3
△G = 0.0 kcal/mol
Target-4
△G = 0.0 kcal/mol
H6
H6
H6
Target-5
△G = − 29.2 kcal/mol
Target-6
△G = − 2.5 kcal/mol
Target-7
△G = − 4.9 kcal/mol
Target-8
△G = − 5.5 kcal/mol
Target-9
△G = − 5.8 kcal/mol


S7 Fig.

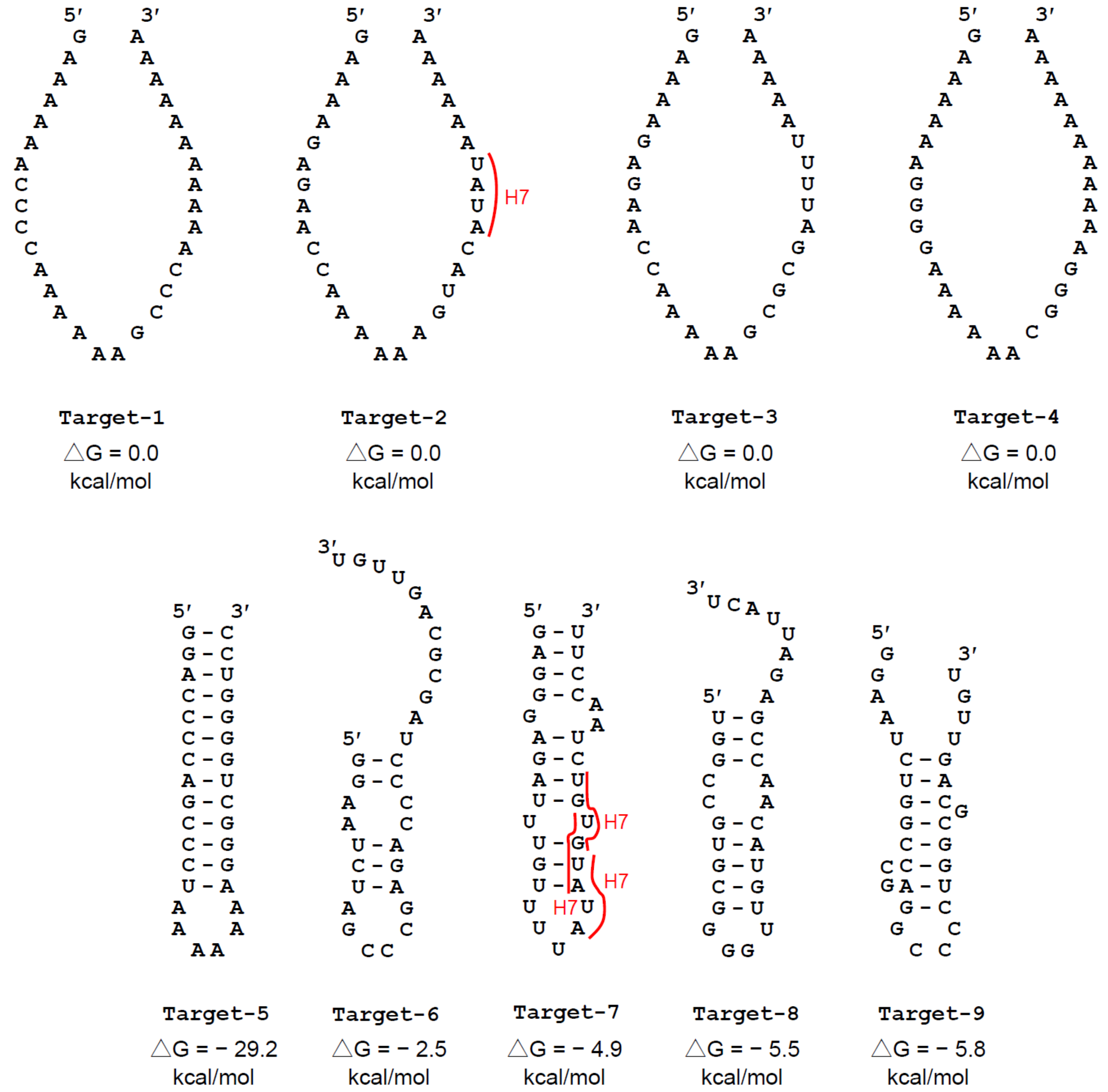
Target-1
△G = 0.0 kcal/mol
Target-2
△G = 0.0 kcal/mol
Target-3
△G = 0.0 kcal/mol
Target-4
△G = 0.0 kcal/mol
Target-5
△G = − 29.2 kcal/mol
Target-6
△G = − 2.5 kcal/mol
Target-7
△G = − 4.9 kcal/mol
Target-8
△G = − 5.5 kcal/mol
Target-9
△G = − 5.8 kcal/mol
H7

S8 Fig.

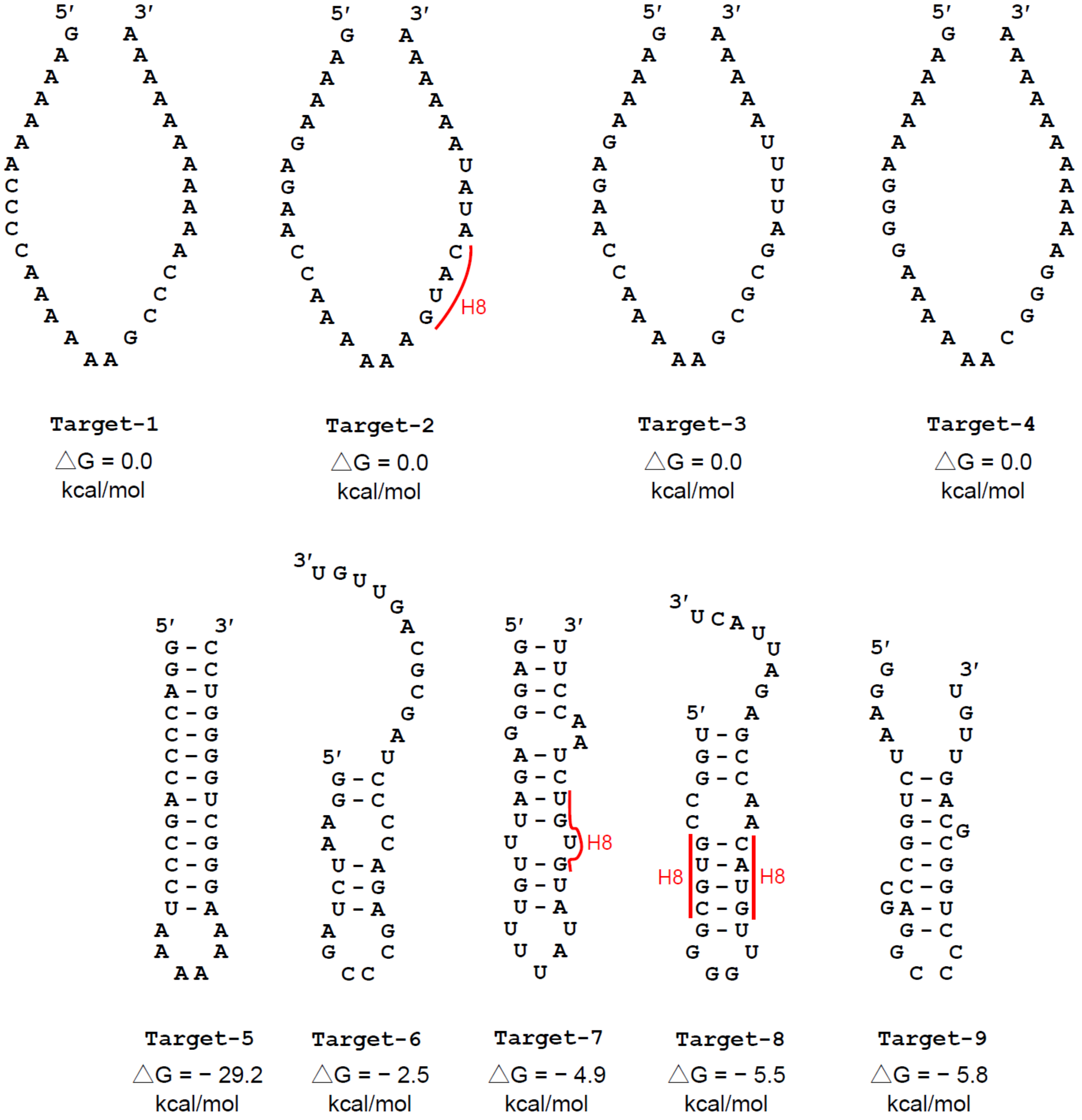
Target-1
△G = 0.0 kcal/mol
Target-2
△G = 0.0 kcal/mol
Target-3
△G = 0.0 kcal/mol
Target-4
△G = 0.0 kcal/mol
Target-5
△G = − 29.2 kcal/mol
Target-6
△G = − 2.5 kcal/mol
Target-7
△G = − 4.9 kcal/mol
Target-8
△G = − 5.5 kcal/mol
Target-9
△G = − 5.8 kcal/mol
H8

S9 Fig.

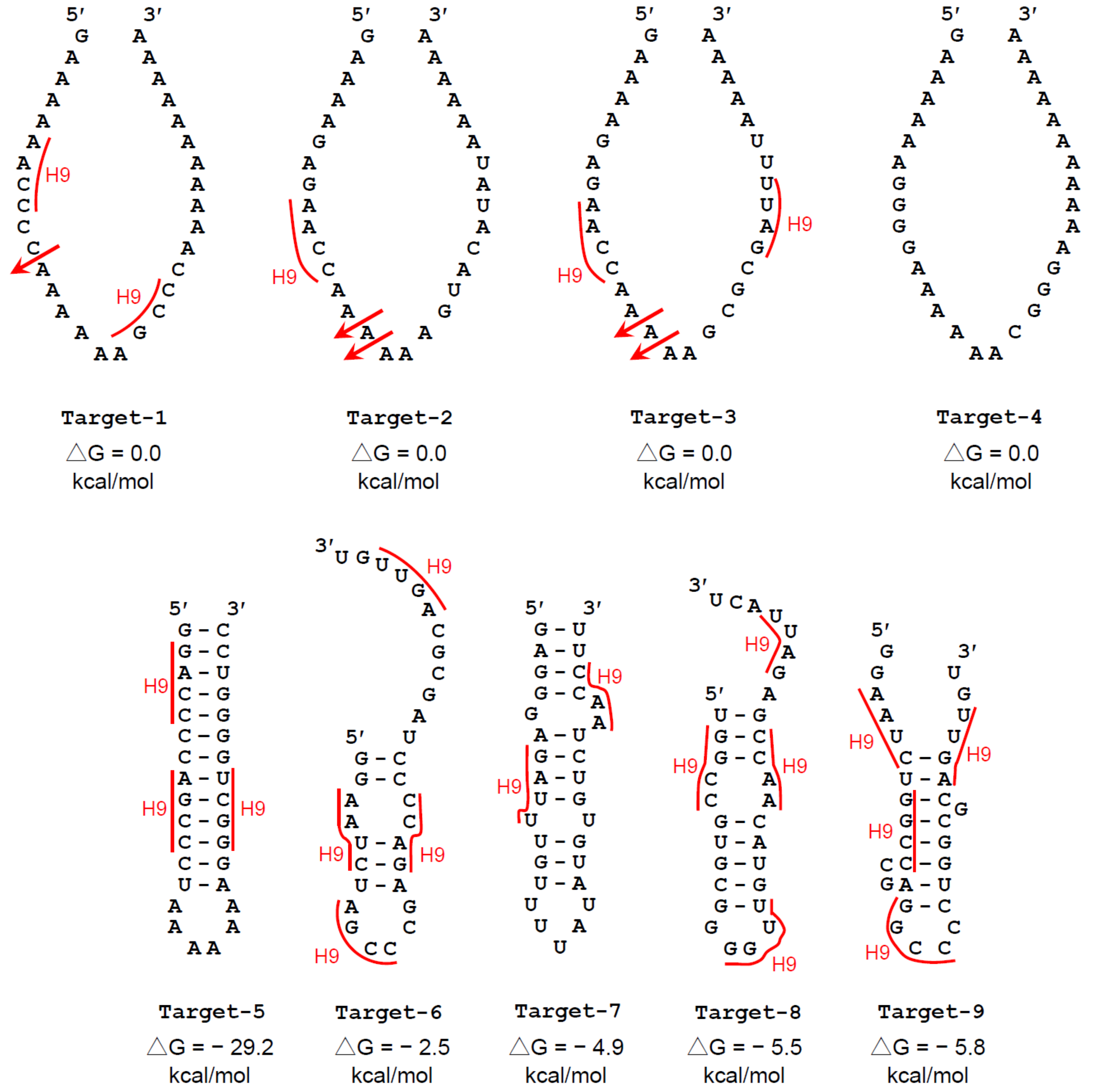
Target-1
△G = 0.0 kcal/mol
Target-2
△G = 0.0 kcal/mol
Target-3
△G = 0.0 kcal/mol
Target-4
△G = 0.0 kcal/mol
Target-5
△G = − 29.2 kcal/mol
Target-6
△G = − 2.5 kcal/mol
Target-7
△G = − 4.9 kcal/mol
Target-8
△G = − 5.5 kcal/mol
Target-9
△G = − 5.8 kcal/mol
H9

S10 Fig.

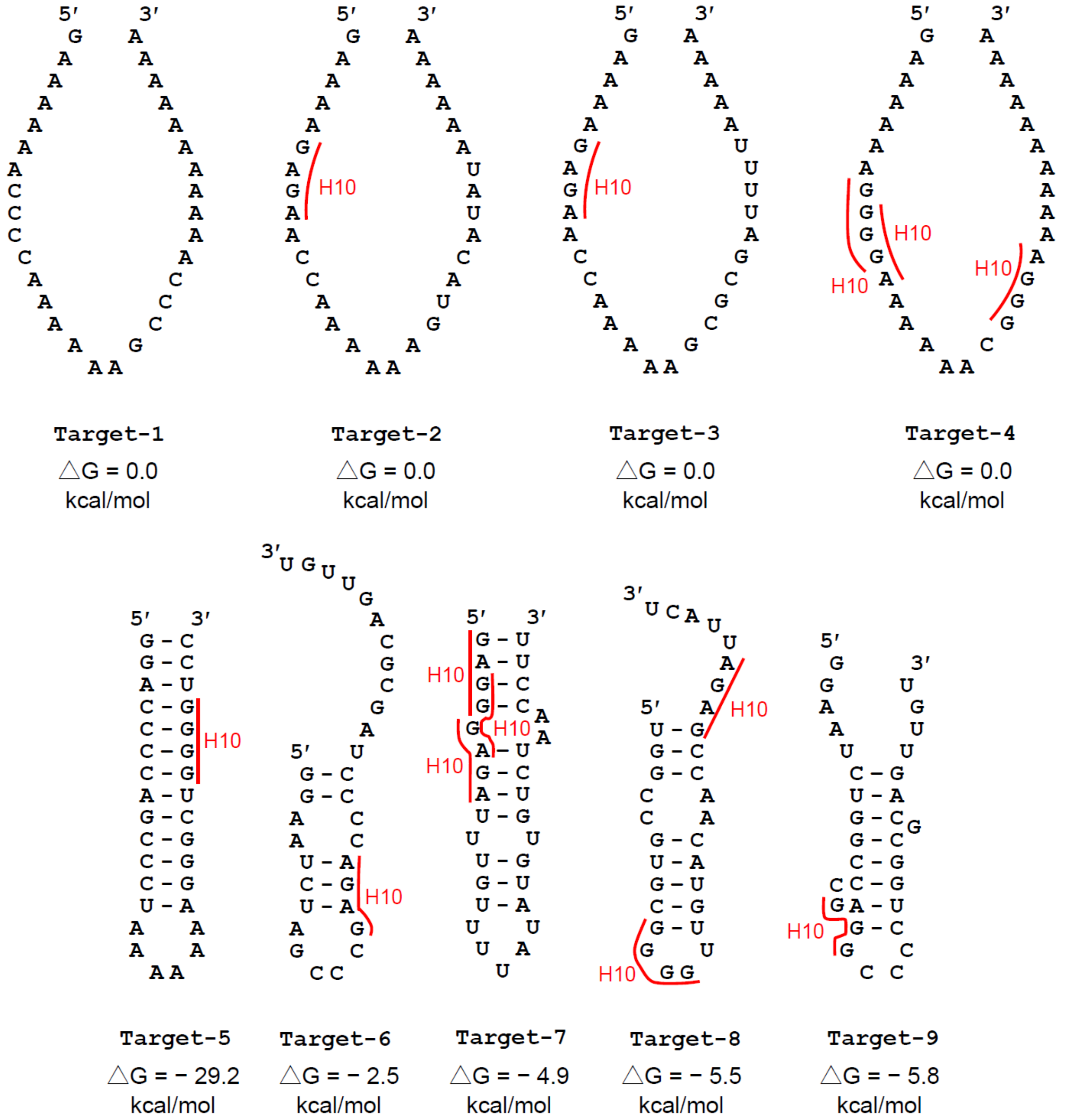

Target-1
△G = 0.0 kcal/mol
Target-2
△G = 0.0 kcal/mol
Target-3
△G = 0.0 kcal/mol
Target-4
△G = 0.0 kcal/mol
Target-5
△G = − 29.2 kcal/mol
Target-6
△G = − 2.5 kcal/mol
Target-7
△G = − 4.9 kcal/mol
Target-8
△G = − 5.5 kcal/mol
Target-9
△G = − 5.8 kcal/mol
H10


S11 Fig.

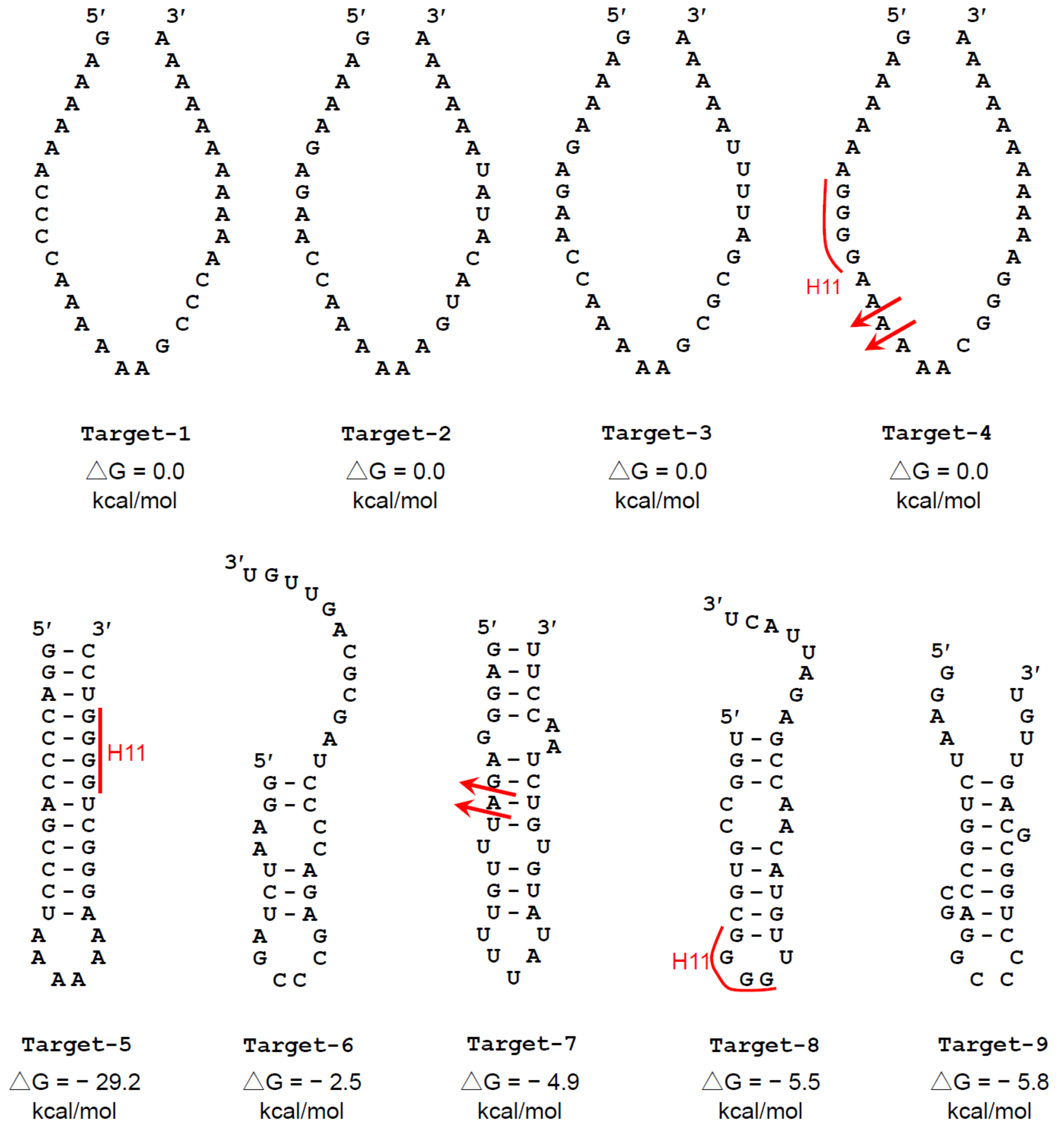

H11
Target-1
△G = 0.0 kcal/mol
Target-2
△G = 0.0 kcal/mol
Target-3
△G = 0.0 kcal/mol
Target-4
△G = 0.0 kcal/mol
Target-5
△G = − 29.2 kcal/mol
Target-6
△G = − 2.5 kcal/mol
Target-7
△G = − 4.9 kcal/mol
Target-8
△G = − 5.5 kcal/mol
Target-9
△G = − 5.8 kcal/mol


S12 Fig.

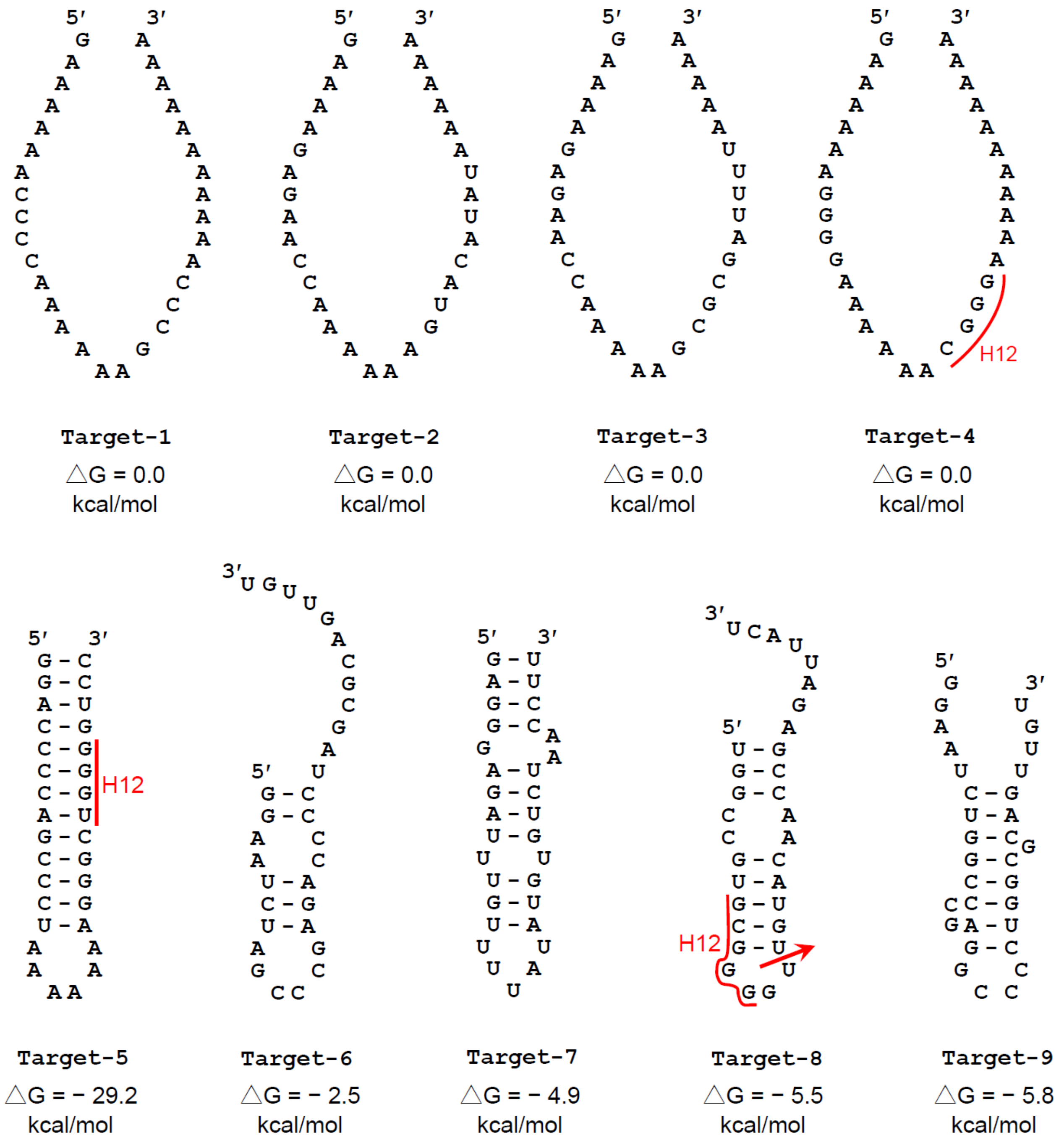
Target-1
△G = 0.0 kcal/mol
Target-2
△G = 0.0 kcal/mol
Target-3
△G = 0.0 kcal/mol
Target-4
△G = 0.0 kcal/mol
H12
Target-5
△G = − 29.2 kcal/mol
H12
Target-6
△G = − 2.5 kcal/mol
Target-7
△G = − 4.9 kcal/mol
Target-8
△G = − 5.5 kcal/mol
H12
Target-9
△G = − 5.8 kcal/mol

S13 Fig.
